# Tertiary lymphoid structures support the development of allergen-specific progenitor CD4+ T cells

**DOI:** 10.1101/2025.10.03.680350

**Authors:** Derek J. Bangs, Brian Hondowicz, Kathleen Abadie, Remi Savard, Madeleine Weiss, Triantafyllia Karakousi, Peter J. Skene, ImmGenT Consortium, Amanda W. Lund, Ananda W. Goldrath, Maximilian Heeg, Marion Pepper

**Affiliations:** Department of Immunology, University of Washington School of Medicine, Seattle, WA, USA; Allen Institute for Immunology, Seattle, WA, USA; Ronald O Perelman Department of Dermatology, NYU Grossman School of Medicine, New York, NY, USA; School of Biological Sciences, Department of Molecular Biology, University of California, San Diego, La Jolla, CA, USA

**Keywords:** TCF1, tissue-resident, CD4+ T cells, tertiary lymphoid structure, iBALT

## Abstract

Tissue-resident memory CD4+ T cells (T_RM_) are key sentinels of the adaptive immune response that provide a rapid, robust inflammatory response upon reactivation in non-lymphoid tissues. While CD4+ T_RM_ are highly protective during reinfections or tumor growth, they are also critical mediators of autoimmunity and allergic disease. Using transcriptional analysis and flow cytometry we profiled the heterogeneity of allergen-specific CD4+ T_RM_ in the lungs following house dust mite exposure and observe two distinct populations of cells: a proinflammatory Th2 lineage and a progenitor TCF1+ lineage that can repopulate the Th2 branch. Confocal microscopy revealed that these two subsets occupied distinct anatomical niches in the inflamed lungs, with Th2 cells localized to the airways while TCF1+ cells localized within pulmonary tertiary lymphoid structures (TLS). Spatial transcriptomics affirmed the TLS as a tissue progenitor niche and highlight the transcriptional progression from progenitor to Th2 cell reflected in the TLS:airway axis. Manipulations to promote or ablate TLS development resulted in increased or decreased TCF1 expression among allergen-specific T cells, respectively. Finally, we identify the PD1 pathway as a critical signal localized to the TLS core and demonstrate that TCF1+ cells in the TLS are responsive to anti-PD1 treatment. Together, these data shape our understanding of tissue CD4+ T cell responses across space and time and highlight TLS as a critical therapeutic target that promotes the propagation of chronic inflammatory diseases.

## Introduction

Tissue-resident memory T cells (T_RM_) are a critical component of the adaptive immune response that provide potent recall responses at the site of exposure in non-lymphoid tissues. While CD4+ T_RM_ are known to have protective roles against a variety of infections and cancer models across tissues (1–4), they can also induce tissue pathology in many settings, including autoimmune diseases or allergic diseases (5–8). While T_RM_ were initially considered a uniform population, recent work has begun to uncover substantial inter- and intratissue heterogeneity that exists within both CD8+ and CD4+ T_RM_ populations (9–12). Developing a more complete understanding of the full spectrum of T_RM_ states will help advance new treatment modalities across a range of human diseases.

A key aspect of tissue immunity that is not well understood is the generation and function of tertiary lymphoid structures (TLS), ectopic lymphoid organs that develop within tissues in settings of chronic inflammation and can function as organizing hubs directly at the affected site. While the exact parameters used to define a TLS can depend on the context, they are typically marked by densely clustered B cells and T cells. Mature TLS can develop features of lymph nodes, including segregated T cell zones, B cell zones marked by follicular dendritic cell (FDC) networks, and high endothelial venules (HEVs) (13). Despite the development of TLS in a variety of inflammatory contexts, including cancers (14, 15), infections (16–18), autoimmunity, (19–22) and asthma (23–25), the contributions of the TLS environment to tissue immunity and T_RM_ differentiation remains unclear.

Asthma is a chronic inflammatory disease characterized by acute, recurrent episodes of pulmonary inflammation that can cause severe clinical disease and even death. Asthma impacts an estimated 25 million people in the U.S., and the prevalence has steadily increased in recent years, similar to many allergic diseases (26). While multiple endotypes of asthma have been described, the most common is allergic asthma, characterized by the production of the type 2 cytokines IL-4, IL-5, and IL-13 following allergen exposure. Treatments for asthma primarily target these acute inflammatory factors (27), and no known cure currently exists. Our lab has previously utilized intranasal exposure to house dust mite extract (HDM) to induce allergic airway inflammation in mice, and we have identified a population of HDM-specific CD4+ T_RM_ that form in the lungs (7). These CD4+ T_RM_ have characteristics of Th2 cells and were critical for the development of disease. However, our understanding of the total diversity of T cell states elicited by HDM exposure and how these heterogeneous populations in the tissue contribute to pathology and sustained chronic inflammation remains incomplete.

Here, we utilized the house dust mite murine model to interrogate both the diversity of CD4+ T_RM_ and how tissue niches in the lungs contribute to T_RM_ fate. Through transcriptional analysis and flow cytometry, we identify both a population of effector Th2 T_RM_ and a precursor population of TCF1+ CD4+ T cells in the lung tissue that can replenish the Th2 pool. These distinct lineages of T_RM_ reside in two different regions of the lung tissue, with Th2 cells in the airways and TCF1+ cells in perivascular TLS. These TLS directly support the development of precursor TCF1+ CD4+ T cells, and spatial transcriptomics reveal that as cells move away from the TLS they adopt a transcriptional program associated with pathologic effector cells. Additionally, we observe that CD4+ T cells in the TLS experience a range of activating and inhibitory signals, with the cells in the core of the TLS experiencing greater inhibitory signals through PD1. Finally, we observe that TCF1+ cells in the TLS are responsive to anti-PD1 checkpoint inhibition, leading to a shift in the antigen-specific population towards the effector lineage. This study demonstrates that allergen-specific T_RM_ can be maintained in two distinct states, one capable of driving acute pathology and one maintaining disease chronicity. We hypothesize that similar precursor-effector relationships exist in other tissues in chronic diseases such as cancer and autoimmunity, and further understanding this relationship could reveal new therapeutic targets.

## Results

### Lung-resident CD4+ T cells induced by house dust mite contain functional heterogeneity

We have previously observed that house dust mite extract (HDM) exposure can induce lung-resident CD4+ T cells that are responsible for driving allergic airway inflammation (7). To gain greater insights into the T cells induced by allergen exposure, we performed CITE-seq on total antigen-experienced T cells (CD3+ CD44+) taken from the mediastinal lymph node and lung parenchyma (demarcated by a lack of intravascular staining) of mice at three different timepoints post-HDM exposure; 5 days after a primary sensitization, 3 days after secondary allergic challenge, and 30 days after the last secondary exposure (Fig 1a). Altogether, single-cell RNA and protein expression data were obtained for over 8000 individual cells, comprising 15 unique clusters across all timepoints and tissues (Fig 1b). We observed a clear division between the cells derived from the draining lymph node and those derived from the lungs (Fig 1c), fitting with previous studies indicating the lungs provide unique immune environments to support diverse differentiation profiles (28). The lung T cell compartment was primarily comprised of CD4+ T cells, highlighting the critical role of CD4+ T cells in this disease model (Fig 1c, Extended Data 1). Examination of T cell responses over time revealed a dramatically evolving immune landscape. Five days post-primary HDM exposure, T cells were primarily confined to the lymph node, and the early wave of T cells entering the lung tissue was relatively homogenous (Fig 1d). In contrast, cells taken three days following secondary HDM challenge were primarily lung-derived and contained substantial phenotypic heterogeneity that was largely maintained in the memory cells isolated at day 30.

**Figure 1:**
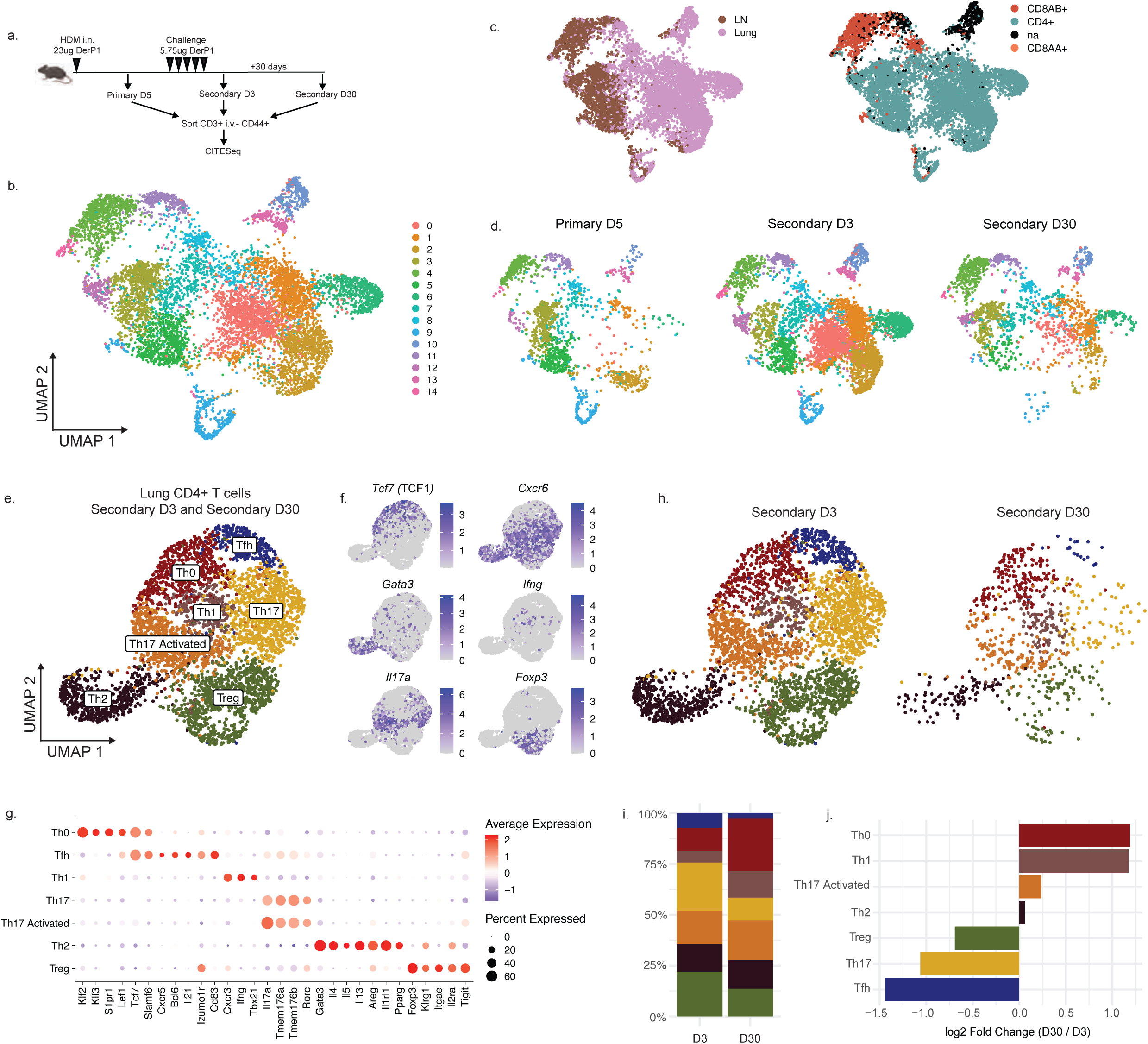
Transcriptional profiling reveals heterogeneous CD4+ T cells following HDM exposure. A) Experimental outline. B) UMAP of all single cells combined across all timepoints (Primary D5, Secondary D3, Secondary D30) and tissues (lungs and lymph nodes). Plots show clusters defined by RNA expression. C) UMAP highlighting tissue of origin or cell type, defined by surface protein expression. D) UMAP showing the breakdown of cells across timepoints. E) UMAP showing reclustered lung CD4+ T cells from Secondary D3 and Secondary D30 timepoints with annotations. F) Plots showing the expression of indicated transcripts. G) Dotplot showing the expression of top differentially expressed genes in indicated clusters. H) UMAP showing lung CD4+ T cells obtained at the indicated timepoint. I) Bar graph showing the relative proportion of cells in each cluster for the indicated timepoint. J) Plot showing the fold change in the proportion of cells at Secondary D30 compared to D3.

To better understand this diversity of cells within the lung tissue, we narrowed our analysis to focus on lung CD4+ T cells in the lung tissue following HDM challenge (Secondary D3 and Secondary D30). Reclustering of lung CD4+ T cells led to the identification of 7 distinct clusters, defined broadly by reciprocal expression of *Tcf7* (encoding TCF1) and *Cxcr6* (Fig 1e,f). The *Cxcr6*-expressing clusters included Tregs (high *Foxp3, Il2ra*), Th1 cells (*Cxcr3, Tbx21, Ifng*), two groups of Th17 cells (*Il17a, Rorc, Tmem176a, Tmem176b*), and a distinct population of Th2 cells marked by *Gata3* and *Il1rl1* (ST2) and expressing high levels of the proinflammatory cytokines *Il4, Il5*, and *Il13* (Fig 1g). The *Tcf7*-expressing clusters included a small Tfh cluster (*Cxcr5, Il21*) and a larger population of cells lacking a clear lineage signature (Th0). Th0 cells identified in the lungs contained small amounts of central memory-associated transcripts (*Sell, Ccr7*, Extended Data 1) but co-expressed high levels of *Lef1*, suggesting they represent a stem-like progenitor subset similar to those described recently in human allergic patients (29). All clusters found at the Secondary D3 timepoint were retained in the tissue at memory (Secondary D30), though we observed differences in the efficiency with which they were preserved (Fig 1h-j). Interestingly, the Th0 cells were the most efficient at retaining a memory population in the tissue, further suggesting they may contain a superior memory capacity. Taken together, these data indicate that HDM challenge leads to the generation of diverse and long-lasting CD4+ T cells in the lungs, including Th0 and Tfh populations defined by high *Tcf7* and a Th2 population defined by expression of *Cxcr6, Gata3,* and *Il1rl1*.

### Derp1-specific CD4+ T cells in the lungs contain functional diversity

We next sought to understand if antigen-specific cells could recapitulate the T cell subset diversity we observe among polyclonal T cells. We have previously utilized a fluorescently-conjugated peptide:MHC tetramer to identify and characterize Derp1:I-A^b^-specific CD4+ T cells in the lungs following HDM administration (7). By using a tetramer constructed with CITE-seq-compatible DNA-tagged streptavidin, we were able to identify a small population of Derp1-specific cells in the sequencing data set. Derp1-specific cells at D3 post-challenge were present in all cell clusters except the cluster containing Tregs, indicating a population of antigen-specific cells can give rise to functionally heterogeneous cells in the tissue (Extended Data 1). To further understand how TCR affinity and antigen specificity overlapped with the functional heterogeneity observed in the lungs, we analyzed the phenotype of expanded clonotypes utilizing the TCR sequences contained in the CITEseq dataset. Interestingly, we observed that each expanded clonotype was found across many different lung CD4+ clusters (Extended Data 1), confirming that cells with the same T cell receptor and therefore affinity for peptide:MHC complexes can produce diverse effector lineages. Additionally, we observed that each expanded clonotype contained some proportion of cells in the Th0 cluster, highlighting the conserved nature of this lineage (Extended Data 1).

To confirm our transcriptional analyses and more thoroughly characterize the functional capacity of Derp1:I-A^b^-specific CD4+ T cells in the lung parenchyma, we utilized flow cytometry combined with a tetramer enrichment strategy (30). Derp1-specific cells reflected the diversity of phenotypes identified by sequencing, with distinct populations of TCF1+ cells and a subset of Th2s expressing CXCR6, ST2, and GATA3 (Fig 2a-c, Extended Data 2). Additional phenotyping of TCF1+ cells confirmed they were not effector cells (CXCR6- and BLIMP1-), Th1 cells (CXCR3-), Tregs (FOXP3-), or TFH (mixed FR4, BCL6-) and instead represented a distinct lineage. Both Th2 cells and TCF1+ cells were able to form memory populations in the tissue (Fig 2d). Similar to the results observed by sequencing, the antigen-specific TCF1+ lineage was enriched over time and became the dominant memory population, while the Th2s had a mild enrichment and the double negative population was largely diminished. Th2 cells expressing ST2 have been indicated to have a pathologic role in allergic asthma (31, 32). Fitting with this, we observed that ST2+ Derp1-specific cells had the highest capacity for production of the type-2 cytokines IL-4, IL-13, and IL-5 following allergen challenge (Extended Data 2) and after a memory restimulation (Fig 2f). This ability of ST2+ cells to selectively produce inflammatory cytokines was reflected in the total CD44+ tetramer(−) population as well (Extended Data 2). Importantly, lymph node Derp1-specific cells contained very few ST2+ cells and had minimal IL-4, IL-13, and IL-5 production (Extended Data 2), highlighting the unique differentiation signals provided by the lung tissue that is required for mature Th2s (28). Together, these data demonstrate that Derp1-specific cells in the lungs form distinct populations of TCF1+ and ST2+ cells early after allergic challenge that form immunological memory and retain distinct functional profiles.

**Figure 2:**
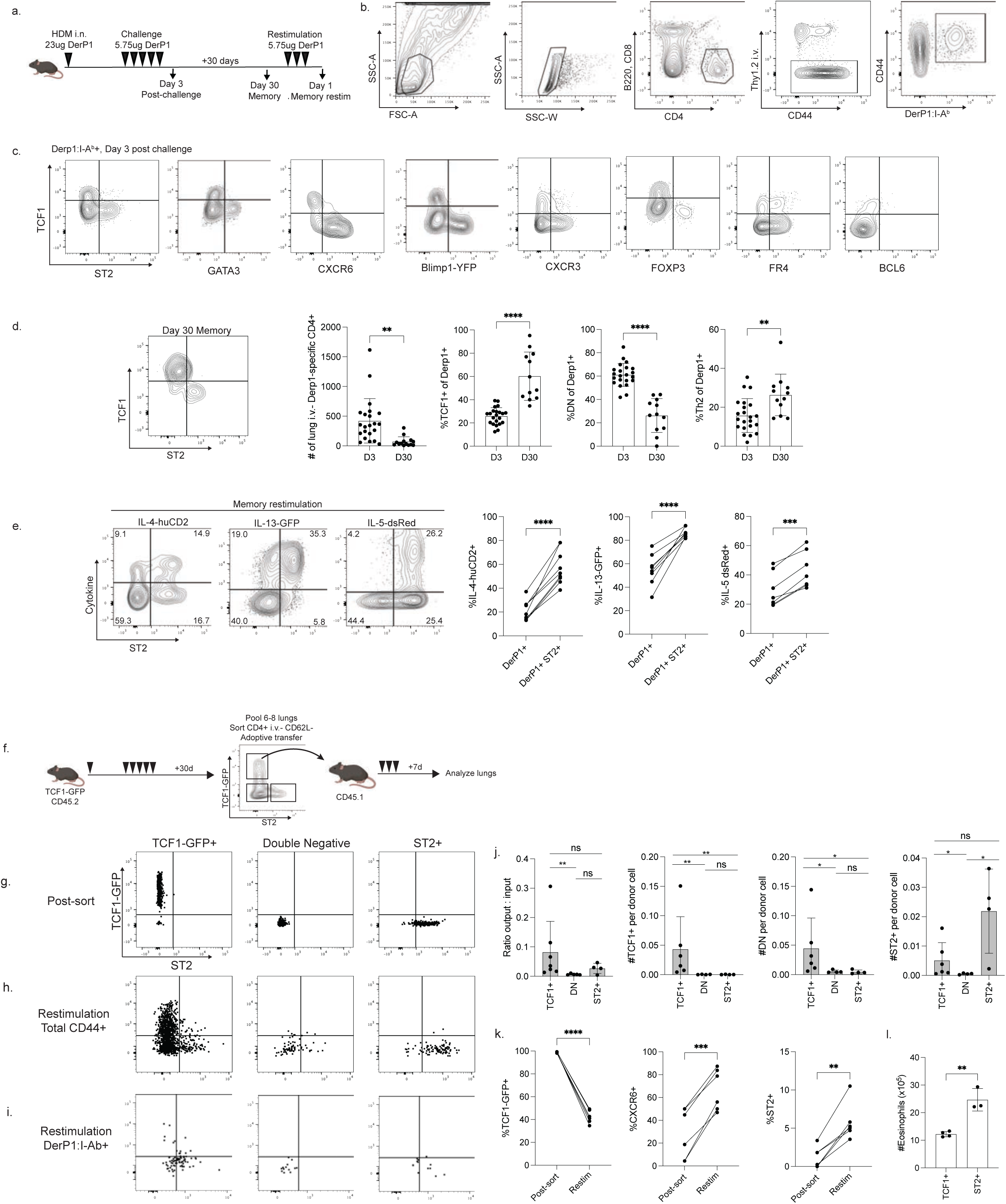
Allergen-specific CD4+ T cells contain proinflammatory Th2s and progenitor TCF1+ cells. A) Outline of experimental design and HDM exposures. B) Representative flow cytometry gating strategy to identify antigen-specific CD4+ T cells. C) Flow cytometry plots, gated as shown in (B), displaying TCF1 staining by the indicated marker on the x-axis. D) Representative flow cytometry plot and summary data showing the numbers and phenotype of Derp1-specific T cells 30 days post-HDM challenge compared to 3 days post-HDM challenge. E) Flow cytometry plots on gated Derp1-specific cells showing expression of ST2 on the x-axis and the indicated cytokine reporter on the y-axis following a restimulation of memory cells, as outlined in (A). Summary data is at right. F) Experimental outline of adoptive transfer. G) Phenotype of sorted cells before transfer. H) Representative flow plots showing the phenotype of transferred CD45.2+ CD45.1-cells following adoptive transfer and restimulation. I) Representative flow plots of Derp1:I-A^b^+ cells among the transferred CD45.2+ CD45.1-cells. J) Summary data showing the ratio of the number of cells of each phenotype recovered in the recipient mouse compared to the number of cells initially transferred for each population. K) Summary data of the phenotype of the transferred TCF1-GFP+ cells. Data show the expression of TCF1-GFP, CXCR6, and ST2 immediately after sorting or following adoptive transfer and restimulation. L) Counts of eosinophils in the lungs of recipient mice given either TCF1-GFP+ cells or ST2+ cells in adoptive transfer. Data in F are combined from three independent experiments and statistical significance was determined with a paired t-test. Data in H-M are combined from three or more independent experiments. Statistical significance was determined using an unpaired t-test in K and M, and a paired t-test in L.

### TCF1+ CD4+ T cells are Th2 precursors and are partially BCL6-dependent

IL-4, IL-13, and IL-5 production has been demonstrated to induce clinical symptoms of asthma, including eosinophilia (33) and airway hypersensitivity (34), indicating a clear role for ST2+ CXCR6+ Th2 cells in driving acute allergic disease. However, we sought to understand how TCF1+ CXCR6-ST2-cells may contribute to disease. Previous studies demonstrated that TCF1+ T cells can have stem-like capabilities across a variety of inflammatory settings, including the ability to generate *de novo* effector populations (35–39). We hypothesized the TCF1+ lineage may function as a progenitor subset for the CXCR6-expressing lineages. To test this, we performed adoptive transfers utilizing TCF1-GFP reporter mice (Fig 2f). Following adoptive transfer, ST2+ cells did not change their identity (Fig 2h), indicating they are likely terminally differentiated. Double negative cells (DN, TCF1-GFP- and ST2-) primarily stayed DN, though some were able to upregulate ST2. In contrast, TCF1+ cells showed plasticity, generating *de novo* DN and ST2+ populations, upregulating CXCR6 and downregulating TCF1 (Fig 2k). Among the transferred cells, we were able to detect rare Derp1:I-A^b^-specific cells (Fig 2i). These cells followed a similar trend, with the DN and ST2+ populations not changing their phenotype but the TCF1+ population showing precursor capacity. In contrast to the total CD4+ population, the transferred antigen-specific TCF1+ cells were more efficient at upregulating ST2, suggesting antigenic stimulation may be critical for promoting terminal differentiation of TCF1+ precursors into Th2s.

In addition to the capacity to generate more terminally differentiated populations, TCF1+ cells were the most efficient at establishing lung residency following adoptive transfer, with the highest ratio of recovered cells:transferred cells (Fig 2j). ST2+ cells were moderately efficient following transfer, while DN cells were highly inefficient. Additionally, all three populations were much more efficient at trafficking to the lungs compared to their relative inability to traffic to the lymph node (Extended Data 3). We also measured eosinophilia in the lungs to determine if the transferred cells were capable of driving inflammation. The mice that received ST2+ cells had elevated lung eosinophilia compared to mice that received TCF1+ cells (Fig 2l), fitting with the greater capacity of ST2+ cells to generate IL-5. Together, these data demonstrate that ST2+ cells are capable of inducing acute inflammation, while TCF1+ cells can function as precursors to ST2+ cells.

TCF1+ BCL6- precursor cells that arise during chronic viral infection have been shown to be dependent on an early BCL6-expressing population for their optimal development (37). To determine the role of cell-intrinsic BCL6, we generated mixed bone marrow chimeras. BCL6 deficient cells had a reduced TCF1+ population of Derp1-specific cells in the lungs following HDM challenge (Extended Data 3), despite not actively expressing BCL6 protein or transcript (Fig 2c, Extended Data 1). Similar results were observed among memory Derp1-specific cells and for total CD44+ cells at both timepoints (Extended Data 3). These results indicate TCF1+ progenitor cells in this system are dependent on BCL6 despite not actively expressing BCL6.

### TCF1+ CD4+ T cells and Th2 cells show distinct localization patterns in the lungs

After observing differences in the expression of chemokine receptors among CD4+ T cells in the lungs, most notably CXCR6, we hypothesized different lineages would acquire different localization patterns. To address this, we performed confocal microscopy on lungs of mice three days post-HDM challenge. We observed CD4+ cells throughout the tissue, with cells in close proximity to the bronchi (EpCAM+), alveoli (podoplanin+), and blood vessels (autofluorescence+ EpCAM-, Extended Data 4). In addition, some CD4+ cells localized with B cells within tertiary lymphoid structures (TLS) that developed following HDM challenge. To track the localization of Th2 cells among these different environments, we imaged the lungs of IL-13 GFP mice. IL-13+ CD4+ cells were found throughout the tissue, with a majority found in the alveolar space and close to bronchi (Fig 3a,h). Interestingly the TLS contained very few IL-13+ cells despite being heavily enriched for CD4+ cells overall. To further confirm the localization of Th2 cells, we tracked IL-5-producing cells using Red5 (IL-5-dsRed) mice. IL-5+ CD4+ cells were also found primarily in the alveolar space and close to bronchi, while comprising a minimal fraction of CD4+ cells in the TLS (Fig 3b,h). To identify the localization of the progenitor subset, we stained for TCF1 in the tissue and observed that the TLS contained a unique and dramatic enrichment for TCF1+ CD4+ cells compared to other tissue sites (Fig 3c,h). TCF1-GFP reporter mice confirmed this result (Fig 3d,h). Together, these data demonstrate that the functional bifurcation between TCF1+ and Th2 cells is reflective of a spatial bifurcation, with TCF1+ cells occupying the TLS and Th2 cells occupying the alveoli and bronchi.

**Figure 3:**
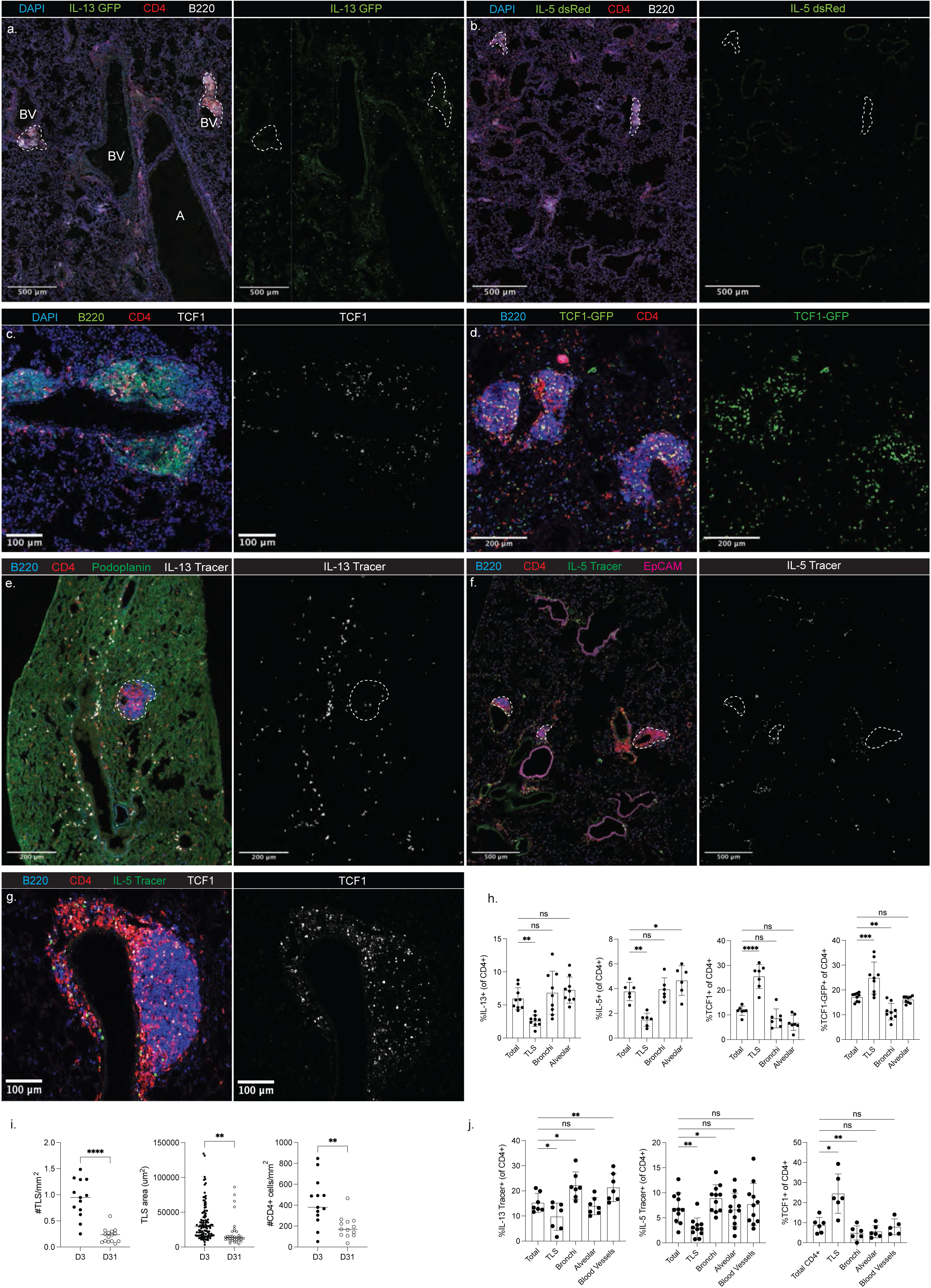
Spatial heterogeneity among lung CD4+ T cells following HDM exposure. Lung tissue sections were generated from mice treated with HDM as outlined in Figure 2A. A,B) Representative images taken of lung sections from IL-13-GFP mice (A) and Red5 mice (B) three days post HDM challenge. C,D) Representative image of TCF1 staining (C) or TCF1-GFP reporter imaging (D). E,F) Representative images of lungs from Yet13Cre-Ai9 (“IL-13 Tracer”) mice (E) and Red5-Ai6 (“IL-5 Tracer) mice (F) 31 days post-HDM exposure. G) Representative image of TCF1 staining in the lungs 31 days post-HDM exposure. H) Summary data from imaging performed at three days post-challenge. I) Summary data showing the density of TLS, size of each individual TLS, and density of CD4+ T cells comparing imaging at three and 31 days post-HDM exposure. J) Summary data from imaging performed at 31 days post-challenge. In H and J, CD4+ cells were categorized as close to TLS (<50um), bronchi (<50um), and blood vessels (<100um, >50um from TLS) or alveolar (>50um from TLS, >50um from bronchi, >50um from blood vessels). Each dot represents data from one image, and images were acquired from three different mice. For each mouse, three different images were acquired from tissue sections generated at least 100um apart from each other. Statistical significance determined using a t-test.

Previous studies describing follicular structures in the lungs have termed these regions inducible Bronchus-Associated Lymphoid Tissue (iBALT) (40). However, the TLS that developed following HDM exposure were only occasionally associated with bronchi, but were consistently formed in the perivascular space. HDM-induced TLS lacked some markers of full maturation, including CD35+ follicular dendritic cells (FDCs) and PNAd+ high endothelial venules (Extended Data 4). Despite the lack of FDCs, some TLS contained large numbers of B cells expressing the transcriptional regulator BCL6, suggesting the TLS were capable of supporting a nascent germinal center response directly within the tissue (Extended Data 4). In contrast to B cells, CD4+ T cells in the TLS expressed very little BCL6 protein, though they also had elevated Ki67 and Nur77, indicating antigenic stimulation (Extended Data 4). Lymphatic vessels can regulate the formation of TLS and the trafficking of T cells with a progenitor phenotype (22, 41). HDM exposure led to an increase in lymphatic vessel density in the lungs, and TLS were markedly enriched for lymphatics compared to other lung regions (Extended Data 4). Together these data demonstrate that HDM exposure leads to the development of perivascular TLS that represent a unique tissue niche for lymphocytes

### Lung-resident CD4+ T cells exhibit spatial heterogeneity at memory

After observing clear spatial heterogeneity among Th2 cells and TCF1+ CD4+ T cells in the lungs following acute HDM exposure, we aimed to determine if this pattern was maintained at memory time points. Interestingly, we observed that TLS were retained at memory but experienced a contraction, with a reduced frequency of TLS in the tissue and reduced average TLS area coinciding with an overall reduction in the density of CD4+ cells (Fig 3i). As a strategy to track memory Th2 cells, we utilized both IL-5 fate mapping mice (Red5-Ai6, “IL-5 Tracer”) and IL-13 fate-mapping mice (Yet13Cre-Ai9, “IL-13 Tracer”). Flow cytometry confirmed these fate-mapping mice efficiently marked cells expressing CXCR6, ST2, and GATA3 (Extended Data 4), allowing us to track memory Th2 cells. Both total IL-13 traced cells and CD4+ IL-13 traced cells were found throughout the lung tissue and enriched around bronchi and blood vessels (Fig 3e,j), fitting with a previous study identifying these regions as important hubs for ILC2s and Th2s (42). Importantly, IL-13 traced CD4+ cells were mostly absent in the TLS. Mirroring the distribution observed in the IL-13 Tracer mice, IL-5 traced cells were found near bronchi and blood vessels, and CD4+ IL-5 traced cells were slightly enriched around bronchi and absent in TLS (Fig 3f,j). Finally, we examined the localization of memory TCF1+ CD4+ cells in the lungs and found that they were primarily found in the TLS (Fig 3g,j). Together, these data clearly demonstrate a distinction in memory niches in the tissue, with the TLS representing long-lived niches for TCF1+ cells, while memory Th2 cells localize to the bronchi and blood vessels.

### Spatial transcriptomics reveal transcriptional changes across lung niches

After observing distinct tissue niches for TCF1+ and Th2 cells through microscopy, we aimed to gain a more comprehensive and quantitative understanding of the tissue environment using spatial transcriptomics (Xenium, 10X Genomics). Lung tissue sections were generated at secondary d3 and secondary d30 timepoints following HDM exposure. Detection of *in situ* transcripts allowed the robust identification of key immune and non-immune cell types throughout the tissue, recapitulating nearly all major cellular lineages (Fig 4a,b, Extended Data 5). Neighborhood identification generated six distinct tissue regions that were classified based on localization and cell composition. This process allowed identification of TLS, bronchi, vessels, adventitia (areas surrounding bronchi and vessels), parenchyma, and the capsule (Fig 4c).

**Figure 4:**
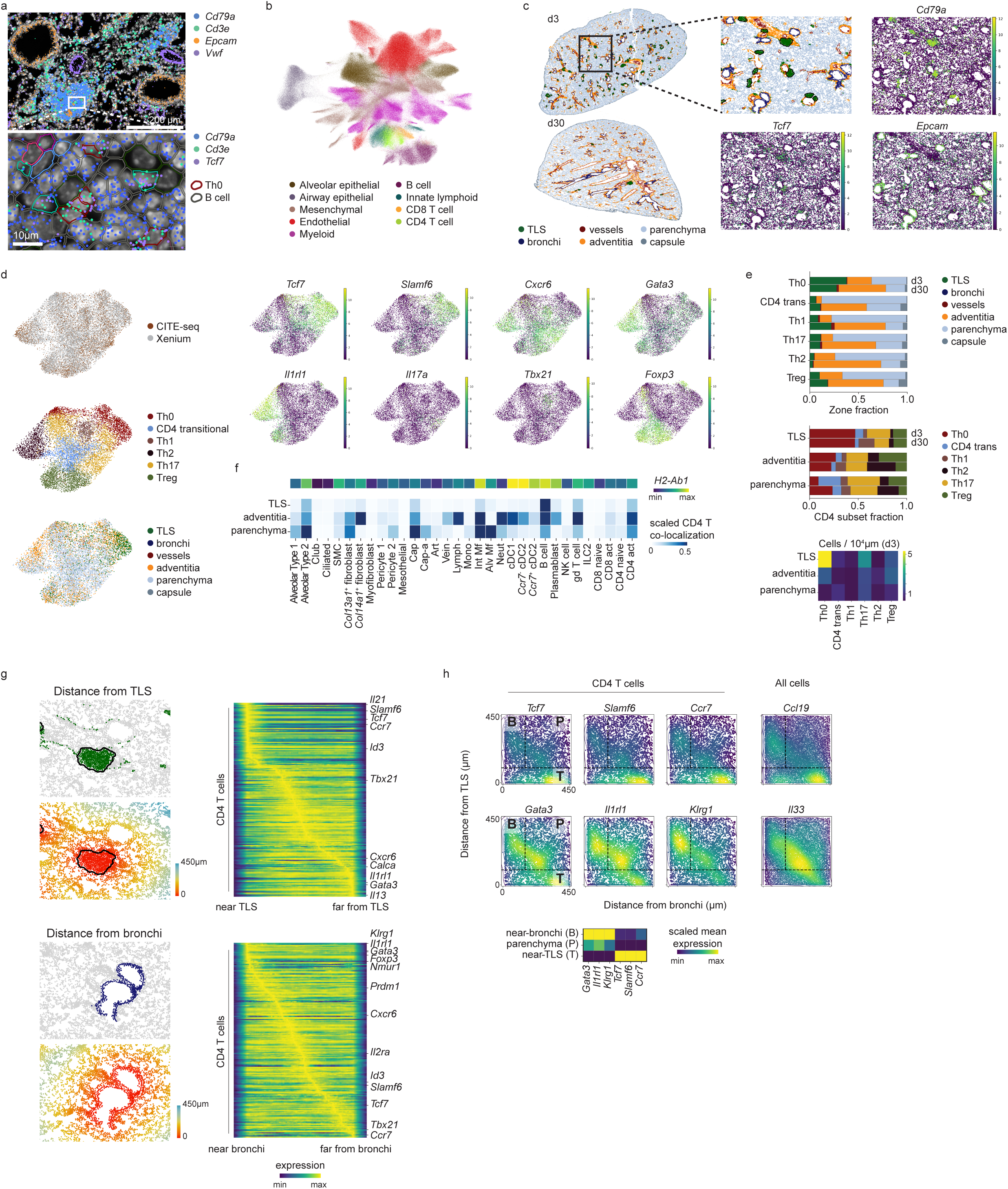
Spatial transcriptomics reveals diverse lung niches. A) Xenium output for day 3 lung with DAPI staining in white, single-cell segmentations outlined, and select transcript locations colored; bottom row shows zoomed view of white box in top row, with cell segmentations colored by cell type. B) UMAP representation of Xenium day 3 and day 30 lung data after scANVI (59) integration with a single-cell RNA-seq reference (56). C) Zones identified using CellCharter (57) on HDM and flu Xenium data at day 3 and day 30; zoomed view of day 3 lung at right shows zones and gene marker expression for B cells (*Cd79a*), Th0 cells (*Tcf7*) and airway epithelial cells (*Epcam*); TLS zone outline in black. D) Activated CD4 T cell subset labeling by integration with CD4+ Lung CITE-seq data: integrated UMAP colored by modality (top left, CITE and Xenium), CD4 subset (middle left, Xenium only), or zone (bottom left, Xenium only); marker gene expression on integrated UMAP (right, Xenium only). E) Activated CD4 T cell subset spatial heterogeneity: zone distribution of all CD4 subsets at day 3 and day 30 (top); composition of CD4 subsets in each major CD4-containing zone (middle); density of CD4 subsets in each major CD4-containing zone by zone tissue area (bottom). F) Squidpy interaction scores at day 3 between activated CD4 T cells and all other cells in each zone (rows are scaled and colorbar maximum is set to 0.5 to visualize low scores), also showing *H2-Ab1* expression per cell type. G) For day 3, distance from TLS (top) or bronchi (bottom) zone, showing convolved gene expression in CD4 T cells with each distance axis. H) Kernel density estimates weighted by gene expression for CD4 T cells at day 3 (left) or all cells (right, sampled to 50,000 cells for visualization) placed on 2D axes of distances to TLS and bronchi zones, with spatial regions close to bronchi (B), in parenchyma (P), or close to TLS (T) indicated; CD4 T cell mean gene expression by region is shown in heatmap below.

After confidently identifying immune niches in the tissue, we next aimed to understand how these neighborhoods overlaid with the diversity of CD4+ T cells we observed. We integrated the single-cell transcriptional data of CD4+ T cells from the Xenium data set with the CITE-Seq data previously generated (Fig 1). This allowed us to recapitulate the functional diversity we previously observed, with clusters of Th0, Th2, Th17, Th1, and Tregs clearly identified, as well as a transitional population (CD4 trans, Fig 4d). Examination of these CD4 subsets across tissue neighborhoods revealed that the TLS was heavily enriched for the Th0 lineage and had a uniquely high density of Th0-polarized cells compared to other lung compartments. In contrast, the adventitia and parenchyma neighborhoods were dominated by the CXCR6+ effector subsets (Fig 4e). Interestingly, all lineages became more heavily localized to the adventitia at d30 compared to d3, fitting with this structure as a critical memory niche in settings of type-2 inflammation. Detection of the cell types spatially co-localized with CD4+ T cells in each region revealed a changing landscape of antigen-presenting cells across the tissue, with B cells being the predominant *H2-Ab1*+ lineage in the TLS while CD4+ T cells in the adventitia and parenchyma contained co-localization with a heterogeneous mix of dendritic cells and macrophages (Fig. 4f).

To investigate the hypothesis that CD4+ T cell heterogeneity is spatially defined across these changing signaling environments, we next examined transcriptional changes along continuous spatial distance axes outward from TLS and bronchi (Fig. 4g). Examination of CD4+ T cell-expressed transcripts revealed that Th0-associated genes (*Tcf7, Slamf6, Ccr7)* were highly expressed proximal to the TLS (Fig 4g), while increasing distance away from the TLS was associated with the acquisition of effector phenotypes (*Tbx21, Cxcr6*). Cells most distal to the TLS expressed markers of functional Th2s (*Gata3, Il1rl1*, *Il13*, *Calca*). Examination of CD4+ transcripts in relation to the bronchi revealed a relationship inverted from the TLS, with *Gata3* and *Il1rl1* proximal to the bronchi and *Tcf7* and *Ccr7* highly distal (Fig 4g). To further interrogate the progenitor:Th2 axis, we examined the expression of individual transcripts in relation to both structures, with the regions proximal to the TLS (T) or bronchi (B) or far from both (parenchymal, P) annotated based on distance thresholds (Fig 4h). Th0-associated genes (*Tcf7, Slamf6, Ccr7*) were specifically enriched in the TLS region, with expression highest far from bronchi but also present close to bronchi, likely reflecting the close proximity of some TLS to bronchi. In contrast, Th2-associated genes (*Gata3*, *Il1rl1, Klrg1*) were found primarily outside the TLS and associated with both the bronchi and parenchyma. High *Il33* expression by all cells in the bronchi and parenchyma regions and high *Ccl19* in the TLS region may implicate differential signaling across space as a factor in generating CD4+ T cell heterogeneity. Together, these results affirm the TLS as a unique progenitor T cell niche in the tissue and identify a TLS:airway axis of CD4+ differentiation associated with the progression from progenitor to terminally differentiated cells.

### Targeting the TLS impacts Derp1-specific TCF1 expression in the lungs

After identifying the TLS as a definitive tissue niche for TCF1+ cells, we sought to manipulate TLS development to alter TCF1 expression. First, we hypothesized that pre-existing TLS from an unrelated exposure, such as a viral infection, may shape the incoming allergen-specific response. To test this, we utilized a model of sterile TLS formation by repeated intratracheal Poly I:C exposures (43). Mice received six doses of either Poly I:C or PBS and were rested for 18 days to allow for the resolution of inflammation and the turnover of innate cell populations (44, 45). Mice were then immunized once with HDM and the Derp1-specific population was analyzed at day 10 (Fig 5a). Fitting with previous findings, repeated Poly I:C exposures led to a significant increase in TLS in the lungs compared to PBS exposure (Fig 5c,d). Poly I:C treatment also led to an increase in the presence of TCF1+ CXCR6-Derp1-specific cells in the lungs (Fig 5b,e). Interestingly, we observed a significant positive correlation between the frequency of lung TLS and the expression of TCF1 among the Derp1-specific cells (Fig 5f), indicating the increase in TCF1+ cells was linked to increased prevalence of TLS.

**Figure 5:**
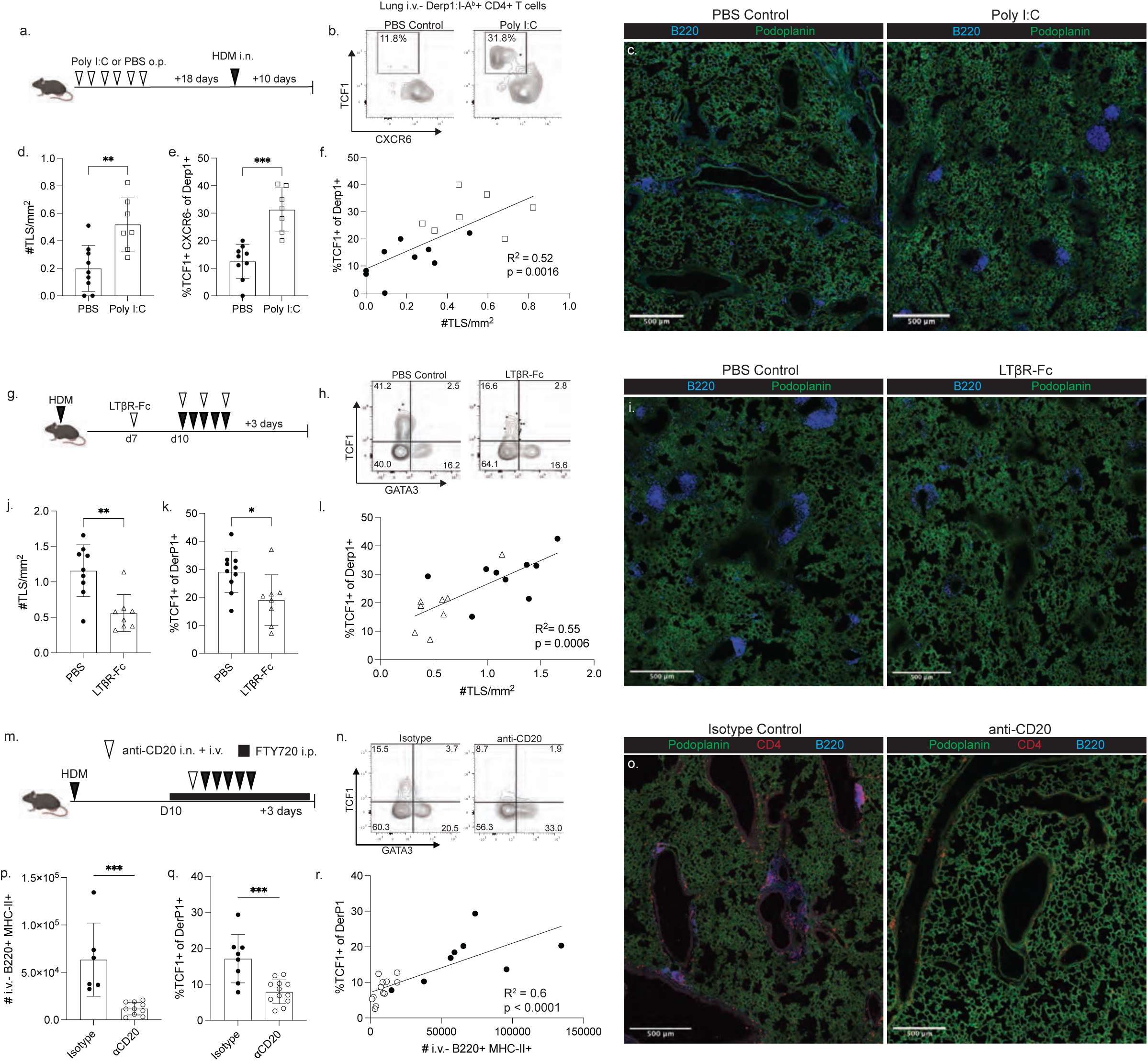
Manipulation of TLS impacts TCF1 expression in lung antigen-specific CD4+ T cells. A) Experimental design of Poly I:C treatment before HDM exposure. B) Representative flow cytometry plot of gated Derp1-specific T cells in the lungs following the indicated treatment. C,D) Representative images of lung tissue sections and summary data of TLS quantification from images. E) Proportion of TCF1+ CXCR6-cells among lung Derp1-specific T cells. F) X-Y plot showing the relationship between TLS density and TCF1 expression for each treated mouse. G) Experimental design of LTβR-Fc treatment during HDM exposure. H) Flow plots of Derp1-specific cells following PBS or LTβR-Fc treatment. I,J) Representative images of lung tissue sections and summary data of TLS quantification from images. K) Proportion of TCF1+ cells among lung Derp1-specific T cells. L) X-Y plot showing the relationship between TLS density and TCF1 expression for each treated mouse. M) Experiential design of B cell depletion. N) Flow plots of Derp1-specific cells following the indicated treatment. O) Representative images of lung tissue sections following anti-CD20 treatment. P) Enumeration of lung i.v.-B cells following the indicated treatment. Q) Proportion of TCF1+ cells among lung Derp1-specific T cells. R) X-Y plot showing the relationship between the number of lung B cells and TCF1 expression for each treated mouse. Data in A-L are combined from 2 independent experiments. Data in M-R are combined from 4 independent experiments. Statistical significance was determined by t test or linear regression (F, L, R).

We next utilized complementary approaches to ablate TLS development during HDM exposure. Lymphotoxin beta receptor (LTβR) signaling has been demonstrated to be critical for the formation of mature TLS, and treatment with soluble LTβR-Fc to antagonize endogenous LTβR signaling has been used to impair TLS formation (46–49). After priming with HDM, we treated mice with LTβR-Fc during the HDM challenge window and assessed the lung and lymph node compartments. LTβR-Fc treatment led to a reduction in lung TLS frequency, a modest reduction in lung parenchymal B cells, and resulted in reduced TCF1 expression among lung Derp1-specific CD4+ T cells (Fig 5h-l, Extended Data 6). Importantly, the acute LTβR-Fc treatment did not significantly alter the lymph node Derp1-specific response or lead to disruptions in the lymph node architecture (Extended Data 6), demonstrating the shift in lung CD4+ T cell phenotype was dependent on changes to the lung environment rather than altered lymph node stimulation. Once again we observed a significant positive correlation between the frequency of TLS and TCF1 expression in the lungs (Fig 5l).

To more thoroughly ablate TLS formation, we utilized an anti-CD20 treatment to deplete B cells. To avoid confounding effects of T:B interactions in the lymph node, we also treated mice with FTY720 to prevent lymphocyte circulation. Importantly, we observed that mice given FTY720 in combination with an isotype control antibody were still capable of generating lung TLS (Fig 5o). Anti-CD20 treatment was highly efficient at depleting B cells in the lung parenchyma and prevented the formation of TLS (Figure 5o,p), resulting in a consistent reduction in the expression of TCF1 expression in Derp1-specific T cells (Fig 5q). In addition, we observed that there was a direct relationship between the number of B cells present in the tissue and the proportion of antigen-specific cells expressing TCF1 (Fig 5r). FTY720 treatment effectively diminished the circulating T cell compartment, and Derp1-specific T cells in the draining lymph node did not change their phenotype or numbers following anti-CD20 treatment (Extended Data 6), definitively demonstrating the changes to the lung Derp1-specific compartment were due to changes in lung signals. Together, these data demonstrate that ablating or promoting TLS formation can reduce or enhance TCF1+ T cells in the tissue.

### TCF1+ CD4+ T cells in TLS are responsive to PD1 checkpoint inhibition

Finally, we sought to identify specific signals in the TLS that could contribute to shaping CD4+ T cell fate. To this end, we calculated the radial distance outward from the TLS centroid for all cells occupying TLS in our D3 spatial transcriptomics dataset, as well as the distance from the TLS outer edge for all cells within 50 µm of a TLS (Figure 6a). We then classified cells in the lowest or highest 50% of the scaled within-TLS radial distance as part of the TLS core region or TLS outer region, respectively, and cells within 50 µm of the TLS edge as the TLS surrounding region. Ordering CD4 T cells by this radial distance (Figure 6b, left) revealed progressive differentiation outward from the TLS center, with cells at the TLS core expressing highest levels of stem-associated genes (*Il7r, Tcf7, Slamf6*), cells in the TLS outer region more highly activated and proliferative (*Mki67, Irf4, Nr4a1*), and cells in the immediate TLS surroundings finally adopting a more differentiated Th2 phenotype (*Cxcr6, Gata3*, *Il1rl1*). Fitting with this, we observed that the TLS contained elevated sources of antigen presentation (*H2-DMb2, H2-Ab1*) and a mix of activating (*Cd86*, *Cd80, Cd40lg, Icosl*) and inhibitory (*Pdcd1lg2, Cd274*) costimulatory signals compared to the area surrounding the TLS (Fig 6b, right).

**Figure 6:**
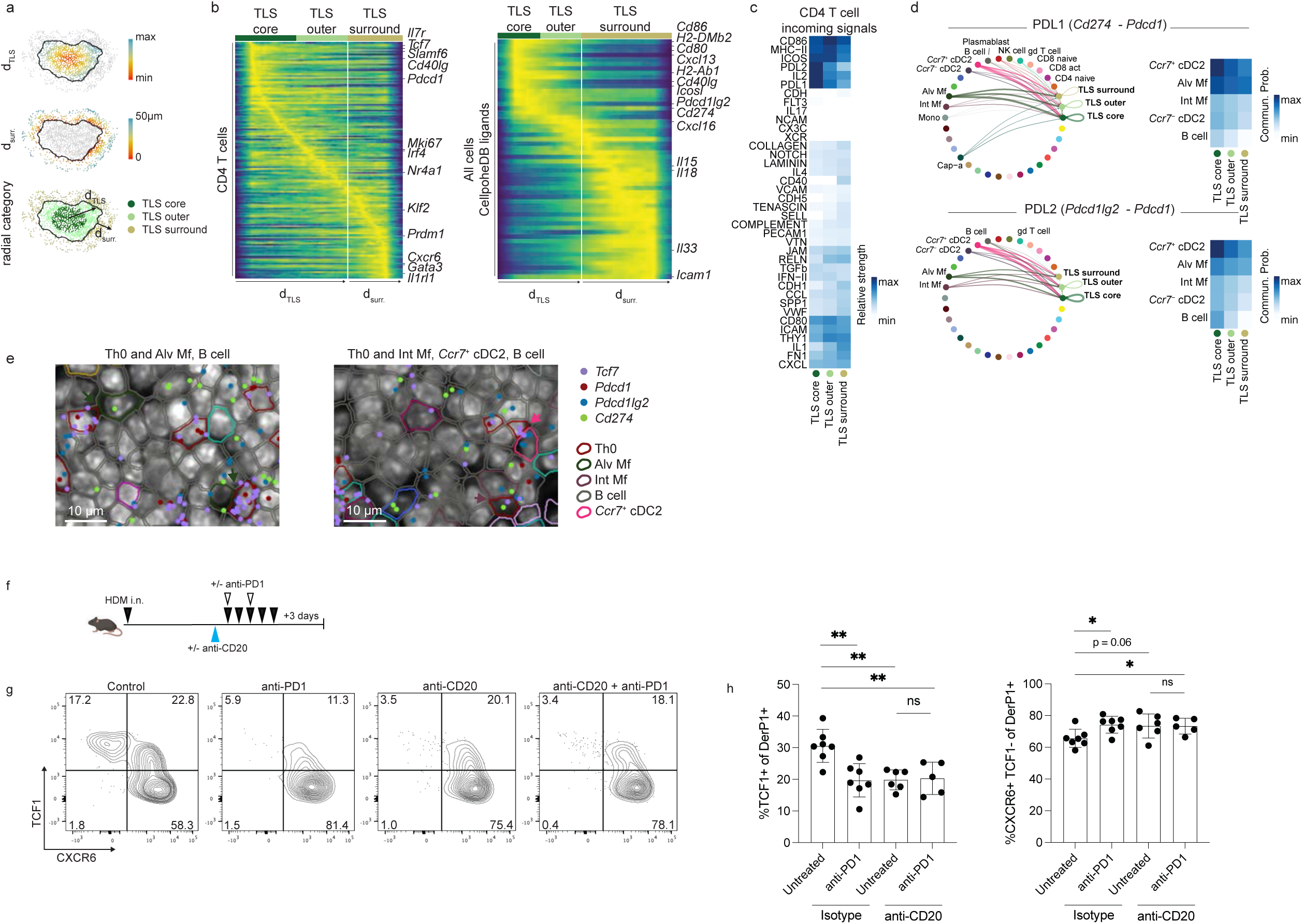
PD1 signaling in the TLS core promotes lung TCF1+ cells. A) For each TLS in the day 3 Xenium sample, the following radial distances were calculated for each cell: radial distance outward from TLS centroid, scaled from minimum to maximum within each TLS (top), radial distance 50 µm from the outer edge of the TLS (middle); cells were subsequently categorized as residing in the TLS core, TLS outer region, and TLS surrounding 50 µm ring. B) Convolved gene expression in CD4 T cells (left) and all cells (right) for cells ordered by distance outward from TLS centroid, with radial categories annotated at top; white vertical line marks barrier between TLS and surroundings. C) Incoming signaling pathways to CD4 T cells in each radial category identified with spatial CellChat (50); signal strength was scaled across all receiver cells (see Extended Data Figure 7) before subsetting to CD4s only and clustering rows. D) Circle plots (left) showing communication probability for sender cells with significant interactions through PDL1 (top) or PDL2 (bottom) with CD4 T cells in the different TLS radial categories, with arc thickness corresponding to communication probability; heatmaps (right) show communication probability for select sender cells to CD4 T cells in different radial categories. E) Snapshots of CD4 Th0 cells in close spatial proximity to potential PD1 signal sender cells in the TLS of the day 3 Xenium sample. F) Experimental outline of PD1 blockade. Prior to HDM challenge, mice were given anti-CD20 or an isotype control, followed by treatment with anti-PD1 or PBS. G) Representative flow cytometry plots gated on Derp1-specific CD4+ T cells in the lung showing TCF1 and CXCR6 expression. H) Summary data showing the phenotype of Derp1-specific CD4+ T cell following the indicated treatment. Data in F-H are combined from three independent experiments. Statistical significance was determined by t test.

To more thoroughly interrogate the signals in the TLS compartments, we performed ligand-receptor interaction analysis on the spatial transcriptomics data (50) to infer the signals CD4+ T cells were receiving in each region as well as the cellular sources. We observed that inhibitory signaling to CD4+ T cells through PDL1 and PDL2 was strongest in the TLS core and decreased progressively moving outward to the TLS outer region and immediate TLS surroundings. This observation that the TLS core contained both high progenitor gene expression in CD4 T cells and high inhibitory signaling led us to hypothesize that inhibitory signals constrain CD4+ T cells and allow them to retain stemness. Fitting with this, we observed that incoming PDL1 and PDL2 signals received by CD4+ T cells were from antigen-presenting cells, including cDC2s, macrophages, and B cells (Fig 6d,e, Extended Data 7).

To directly test if inhibitory signals were critical for the retention of TCF1+ cells in the lung tissue, we treated mice with an anti-PD1 blocking antibody to prevent checkpoint inhibitory signals during HDM challenge (Fig 6f). Derp1-specific cells had robust PD-1 expression that was completely saturated by blocking antibody treatment (Extended Data 7). Anti-PD1 treatment led to a reduction in TCF1 expression and an enrichment for CXCR6+ TCF1-cells among Derp1-specific T cells in the tissue (Fig 6g,h). To see if the changes induced by anti-PD1 treatment were dependent on the development of TLS, we treated mice with anti-CD20 to disrupt TLS formation prior to anti-PD1 treatment. Anti-CD20 treatment effectively depleted lung B cells (Extended Data 7) and reduced TCF1 expression in the tissue (Fig 6g,h). However, anti-PD1 treatment in mice following B cell depletion did not impact the Derp1-specific T cell compartment, demonstrating the effects of anti-PD1 were dependent on the TLS. Together, these data demonstrate that the TLS contains a core region defined by robust PD1 signaling that retains TCF1+ cells that can be disrupted through antibody blockade.

## Discussion

Our results identify a key population of lung TCF1+ CD4+ T cells that occupy tertiary lymphoid structures and can propagate an immune response to allergic exposure. We and others have previously interrogated the development and function of Th2 T_RM_ that are induced by exposure to HDM. Here, we expand our understanding of the tissue CD4+ T cell compartment. We find that lung Th2 and TCF1+ CD4+ T cells have distinct phenotypic profiles, functional capacities, and tissue localization, and that both cells have the capacity to contribute to diseases. While targeting the downstream effector mechanisms of Th2 cells can provide relief from allergic symptoms, therapies targeting TLS or TCF1+ antigen-specific cells may provide novel long-lasting cures for disease.

Our results join other recent studies that have identified functionally heterogeneous populations of lung-resident CD4+ T cells following influenza infection (11,12). These studies described two populations of cells: a proinflammatory Th1 population capable of IFNg production, and a FR4+ resident helper (TRH) population that provides help to B cells and CD8+ T cells. This functional bifurcation in the antiviral immune response directly mirrors the ST2+ and TCF1+ populations we investigate here. The TCF1+ cells that arise following HDM exposure share many similarities with TRH cells, including co-localization with B cells, dependency on B cells for their optimal generation, and dependency on BCL6. However, our results also reveal key differences, including a lack of BCL6, CXCR5, and FR4 expression. Our results support a model proposed by recent work investigating CD4+ responses during chronic LCMV infection, where both a TCF1+ BCL6^hi^ TFH and a TCF1+ BCL6^lo^ progenitor population are formed, with both populations showing a dependency on early BCL6 (37). One possible explanation for the lack of a more robust TRH population in this model would be the lack of fully developed TLS. Influenza infection leads to more widespread and long-lasting inflammation, possibly resulting in a greater capacity for the tissue to support a full-fledged germinal center response and leading to the subsequent development of TRH. Providing additional rounds of HDM stimulation may induce further TLS differentiation and bridge the gap between these two models.

TCF1+ CD8+ and CD4+ T cells have been identified in a variety of settings, including chronic infections, autoimmunity, and cancer. Here, we demonstrate these cells also play a key role in propagating allergic responses. While the role of CD8+ TCF1+ cells has been extensively studied, our understanding of the factors regulating TCF1+ CD4+ T cells is less complete. The ability of CD4+ T cells to differentiate into distinct effector lineages (i.e. Th1, Th2, Th17, etc) adds an additional layer of complexity onto our understanding. This raises the question of if a single TCF1+ pool of cells exists that can give rise to all possible effector lineages, or if TCF1+ cells are already polarized towards a particular T helper lineage. Our results utilizing TCR sequence analysis with transcriptional profiling suggest the former is true, as we observe individual clones found within the TCF1+ pool are found across all helper lineages that arise after HDM exposure. However, additional experiments to formally test this hypothesis are needed to define the exact capacity of these cells.

This study advances our understanding of the role of tertiary lymphoid structures as key players in shaping the tissue immune landscape. While TLS have long been observed to arise in a range of inflammatory conditions across many tissues (13–26), their capacity to contribute to disease and the mechanistic details of that contribution remains unclear. Here, we show TLS can serve as a local reservoir of TCF1+ CD4+ progenitor cells, providing a tissue niche for these cells that positions them for rapid reseeding of the more terminally differentiated lineages. Importantly, we show that preexisting TLS can shape the incoming allergen-specific response, suggesting an unrelated inflammatory event (i.e. respiratory viral infection) can shape the allergic response in a manner previously not appreciated. Our results also highlight the significant role of PD1 signaling in maintaining TCF1+ cells in the tissue, highlighting the core of the TLS provide a critical inhibitory niche otherwise absent in the tissue. This feature of TLS may operate in combination with other mechanisms to promote TCF1+ cells in the tissue, such as continuous recruitment of TCF1+ cells from circulation or the retention of cells through chemotactic signals. Future experiments will be necessary to more thoroughly address these possibilities.

The findings related to TLS function and TCF1+ CD4+ T cell fate have clear implications for future treatment options for allergic diseases but also extends to other disease models driven by CD4+ T cells where TLS are present, including inflammatory bowel disease (IBD, 20-22, 51), autoimmune vasculitis (19), atherosclerosis (52), and inflammatory skin conditions (53). Developing methods to ablate or prevent the development of TLS may help blunt tissue immune responses and thus reduce disease. We show here that treatment with anti-CD20 efficiently cleared TLS from the lungs and led to a reduction in TCF1 expression. Rituximab, a chimeric anti-CD20 monoclonal, has been used for therapeutic purposes to treat cancers and, more recently, autoimmune disorders. Interestingly, Rituximab treatment has been reported to improve clinical asthma, despite not effectively reducing serum IgE levels (54–56). Our results suggest a possible mechanism of this treatment by offering an antibody-independent function of B cells as critical components of the TLS, and this data supports the potential of Rituximab as a treatment for chronic allergic diseases.

Our findings on the role of TLS also have potential implications in tumor immunotherapy. TCF1+ T cells are widely reported to be associated with better clinical outcomes in cancer patients, including a higher responsiveness to checkpoint inhibition (57). TCF1+ cells are known to be the primary cell type responsive to anti-PD-1 checkpoint blockade, though the relevant TCF1+ cells have been reported to come from the draining lymph node (58). In parallel, the presence of TLS has been recently associated with improved clinical outcomes across a range of cancer types (59). Our results link these two observations, suggesting the TLS forms a local population of PD1 checkpoint-responsive TCF1+ cells that can provide more robust anti-tumor responses directly at the tumor site. These findings argue that new methods for targeting TLS formation or disruption should be a future therapeutic priority.

## Methods

### Mice

C57BL/6, B6.SJL-Ptprc^a^Pepc^b^/BoyJ (CD45.1), B6(Cg)-Tcf7^tm1Hhx^/J (TCF1-GFP), B6(C)-Il5^tm1.1(icre)Lky^/J (Red5), B6.Cg-Tg(Prdm1-EYFP)1Mnz/J (Blimp1-YFP),, C57BL/6-Tg(Nr4a1-EGFP/cre)820Khog/J (Nur77-GFP), B6.Cg-Tg(Cd4-cre)1Cwi/BfluJ (CD4-Cre), C.129S4(B6)-Il13^tm1(YFP/cre)Lky^/J (YetCre-13), B6.Cg-Gt(ROSA)26Sor^tm6(CAG-ZsGreen1)Hze^/J (Ai6), and B6.Cg-Gt(ROSA)26Sor^tm9(CAG-TdTomato)Hze^/J (Ai9) mice were purchased from the Jackson Laboratory. Bcl6^flox^ mice were provided by Dr. Alexander Dent (Indiana University). C57BL/6 human CD2 IL-4 reporter mice (KN2) were provided by Dr. Markus Mohrs (Trudeau Institute). C57BL/6 IL-13 eGFP reporter mice were provided by Drs. Andrew McKenzie (MRC) and Michael Holtzman (Washington University). Prox1-CreER^T2^ mice were provided by V.H. Engelhard. All mice were housed, maintained, and bred under specific-pathogen-free conditions at the University of Washington, and all experiments were performed in accordance with the University of Washington Institutional Care and Use Committee guidelines.

### In vivo treatments

Whole house dust mite antigen from *Dermatophagoides pteronyssinus* (HDM, Greer Laboratories) was resuspended in PBS. Mice were anesthetized with ketamine/xylazine and given a dose of HDM at a concentration to provide 23*μ*g of Derp1 protein intranasally for the primary immunization. Mice were subsequently given challenge doses of 5.75*μ*g Derp1 on days 10-14. For memory HDM restimulation, mice were treated with three additional challenge doses of 5.75*μ*g Derp1 following a 31 day rest. For Poly I:C administration, mice were anesthetized with isoflurane and given 40*μ*g Poly I:C (Poly I:C HMW, Invivogen) and 50*μ*g ovalbumin (OVA protein, Invivogen) intratracheally twice a week for three weeks. Mice receiving LTβR-Fc (provided by Saiyu Hang, Genentech) were given 100*μ*g i.p. For B cell depletions, mice were treated with anti-CD20 (clone MB20-11, BioXCell) or an isotype control (control IgG2c, BioXCell) at a combination dose of 250*μ*g i.v. and 100*μ*g i.n. For FTY720 treatment, mice were given 25*μ*g of FTY720 (Enzo Life Science) i.p. For tamoxifen administration, mice were given 1mg of tamoxifen (Sigma) in corn oil i.p. every other day for the duration of treatment. To block PD1, mice were treated with anti-PD1 (clone 29F.1A12, BioXCell) at a combination dose of 250*μ*g i.v. and 100*μ*g i.n on the indicated days.

### Isolation of Single Cell Suspensions

Mice were injected i.v. with 1*μ*g of anti-Thy1.2 or 5*μ*g of anti-CD45 to label circulating T cell or leukocytes, respectively (59). Approximately three minutes after, mice were euthanized via CO_2_ asphyxiation and the lungs were harvested. The lung tissue was placed into gentleMACS C Tubes (Miltenyi Biotec) with RPMI 1640 Medium (Gibco) containing 70*μ*g/mL Liberase ™ (Roche) and 10mM aminoguanidine (Sigma-Aldrich). The tissue was then dissociated on a gentleMACS Dissociator (Miltenyi Biotec) and incubated at 37°C for 30 minutes. Following a final dissociation step, the tissue suspension was filter through a 70*μ*m filter and washed with RPMI 1640 containing 10% fetal calf serum.

### Enrichment of antigen-specific cells and flow cytometry

Single-cell suspensions were stained with Derp1_117-127_ (CQIYPPNVNKI) I-A^b^ tetramer conjugated to APC at room temperature in the dark for one hour. Cells were then washed and stained with 25*μ*L anti-APC microbeads (Miltenyi Biotec) on ice for 30 minutes. After incubation, cells were washed and enriched over a magnetic LS column (Miltenyi Biotec) as previously described (Moon et al., 2007, Legoux and Moon 2012). The tetramer enriched fraction and the non-enriched “flow through” fraction were then stained with surface antibodies on ice for 30 minutes. When applicable, cells were subsequently fixed and permeabilized with the eBioscience Foxp3/Transcription Factor Staining Buffer Set (Invitrogen) according to the manufacturer’s instructions. Transcription factor staining was performed in 1X Permeabilization Buffer at 4°C overnight. Samples were run on a FACSymphony cytometer (BD) and analyzed using FlowJo software (Treestar).

### Bone marrow chimeras

Bone marrow cells were collected from the tibia, femur, humerus, and sternum and labeled with anti-Thy1.2 (30-H12, eBioscience) and anti-NK1.1 (PK136, eBioscience). Cells were resuspended and incubated with low toxicity rabbit complement (Cedarlane Laboratories). After complement lysis, cells were washed with media containing 10% fetal calf serum. Recipient mice were lethally irradiated (1000 rads) and injected intravenously with 5×10^6^ total bone marrow cells with congenically distinct WT cells mixed 1:1 with CD4Cre Bcl6^flox/flox^ cells mixed in equal portions and treated with antibiotic (Enrofloxacin)-treated water for 8 weeks. Prior to house dust mite exposure, chimerism was determined by analyzing the ratio of CD45.1 and CD45.2 expression among cells in the blood.

### Preparation of lung tissue for immunofluorescence

For preparation of lung tissue, mice were euthanized and the lungs were gently inflated with 3mL of 1% paraformaldehyde in PBS (Cytofix, BD Biosciences) injected into the trachea. The trachea was then tied off with a suture string, and the lungs were removed and incubated in 1% paraformaldehyde at 4°C overnight with agitation. The tissue was subsequently washed twice with PBS, incubated at 4°C overnight in a 15% sucrose solution, then incubated at 4°C overnight in a 30% sucrose solution. Lung tissue was then embedded in O.C.T. (Fisher Scientific) and frozen in a slurry of dry ice and 200 proof ethanol.

### Lung Immunofluorescence

Following preparation and freezing, lung tissue was cut into 10*μ*m sections using a cryostat (Leica Biosystems). Sections were mounted on SuperFrost Plus microscope slides (Fisher Scientific) and allowed to air dry for 30 minutes prior to staining procedure. For staining, lungs were rehydrated in PBS for 10 minutes, fixed in 1% paraformaldehyde (Cytofix, BD Biosciences) for 5 minutes, then permeabilized in 0.5% Triton-X for 15 minutes. Sections were blocked with CAS-Block Histochemical Reagent (ThermoFisher) for at least 1 hour at 4°C in a humidified chamber. Sections were stained with antibodies in CAS-Block overnight at 4°C in a humidified chamber. Following staining, sections were washed with PBS, then mounted with Fluoromount-G (ThermoFisher). Confocal images were acquired with a Nikon C2 laser scanning confocal microscope using a 20X objective lens. Analysis of images was performed using ImageJ and Imaris (Oxford Instruments). For analysis of CD4+ T cell phenotypes in Imaris, surfaces were generated using CD4 staining. Additional surfaces were generated for structural features, data was exported for CD4 surfaces as a .CSV file, and expression of fluorescence in the CD4s surfaces classified based on their distance to other lung structures was measured in FlowJo software (Treestar).

### Single-cell RNA, TCR and Totalseq C-Sequencing - Sample Preparation

#### Flow cytometry and HT antibody staining

Single-cell lymphocyte suspensions were isolated as described above and stained with anti-CD45, anti-CD3e, and anti-CD44 AF700 and different TotalSeq-C Anti-Mouse Hashtags (TotalSeq-C Anti-Mouse Hashtags 1-10 Biolegend #155861-155879, in staining buffer (PBS, 2% FCS) for 30min at 4°C in the dark. Standard spleen cell suspensions were stained under identical conditions but included anti-CD45-FITC to differentiate them from other samples when pooled. Each sample was labeled with a unique hashtag, enabling the downstream assignment of individual cell (10x cell barcode) to their respective source samples (60).

##### Cell sorting

For each sample, approximately 50,000 DAPI^−^ CD45^+^ CD3^+^ CD44+ T cells were sorted on the FACSAria III and pooled in a single collection tube at 4C. DAPI was added just before the sort.

##### ImmGenT TotalSeq-C custom mouse panel staining

The pooled single cell suspension was stained with the ImmGenT TotalSeq-C custom mouse panel, containing 128 antibodies (Biolegend Part no. 900004815) and FcBlock (Bio X Cell #BE0307). Because 500,000 cells are required for staining, unstained splenocytes were spiked in to reach a total of 500,000 cells.

##### Second sort

Cells were subsequently sorted a second time with the addition of DAPI to select for live CD45^+^ CD3^+^ T cells. 40,000 CD45^−^ APC^+^ sample cells and 10,000 CD45^−^ FITC^+^ spleen standard cells were sorted into a single collection tube.

### Single-cell RNA, TCR and TotalseqC-Sequencing - Library Preparation

Cell encapsulation and cDNA library. Single-cell RNA sequencing was performed using the 10x Genomics 5’ v2 platform with Feature Barcoding for Cell Surface Protein and Immune Receptor Mapping, adhering to the manufacturer’s guidelines (CG000330). After cell encapsulation with the Chromium Controller, reverse transcription and PCR amplification were performed in the emulsion. From the amplified cDNA library, smaller fragments containing TotalSeq-C-derived cDNA were separated for Feature Barcode library construction, while larger fragments containing transcript-derived cDNA were preserved for TCR and Gene Expression library generation. Library sizes for both cDNA fractions were evaluated using the Agilent Bioanalyzer 2100 High Sensitivity DNA assay and quantified with a Qubit dsDNA HS Assay kit on a Qubit 4.0 Fluorometer.

#### RNA library construction

After enzymatic fragmentation and size selection of the cDNA, the library was ligated to an Illumina R2 sequence and indexed using unique Dual Index TT set A index sequences, TT-H6 and TT-H7, through an index PCR consisting of 14 cycles.

#### TCRαβ library construction

TCR cDNA was amplified from the cDNA using nested PCR following the manufacturer’s protocol. The resulting TCR libraries underwent enzymatic fragmentation, ligation, and indexing with unique Dual Index TT set A (10x part no. 3000431), index sequences, TT-A9 and TT-A10, through an index PCR performed for 8 cycles.

#### TotalSeq-C library construction

Totalseq-C derived cDNA was processed into the Feature Barcode libraries following the manufacturer’s protocol. The library was indexed with a unique dual index TN set A (10x part no. 3000510) index sequence, and TN-C8 and TN-C9 were indexed with an index PCR (8 cycles).

#### Sequencing

The three libraries were pooled based on molarity in the following proportions: 47.5% RNA, 47.5% Feature Barcode, and 5% TCR. The pooled libraries were sequenced on an Illumina NovaSeq S2 platform (100 cycles) using the 10x Genomics specifications: 26 cycles for Read 1, 10 cycles for Index 1, 10 cycles for Index 2, and 90 cycles for Read 2.

### Single-cell RNA, TCR and TotalseqC-sequencing - Data processing

#### Code

Code is available on https://github.com/immgen/immgen_t_git/

#### Count matrices

Gene and TotalSeq-C antibody (surface protein panel and hashtags) counts were obtained by aligning reads to the mm10 (GRCm38) mouse genome using the M25 (GRCm38.p6) Gencode annotation and the DNA barcodes for the TotalSeq-C panel. Alignment was performed with CellRanger software (v7.1.0, 10x Genomics) using default parameters. Cells were identified and separated from droplets with high RNA and TotalSeq-C counts by determining inflection points on the total count curve, using the barcodeRanks function from the DropletUtils package.

#### Sample demultiplexing

Sample demultiplexing was performed using hashtag counts and the HTODemux function from the Seurat package (Seurat v4.1). Doublets (droplets containing two hashtags) were excluded, and cells were assigned to the hashtag with the highest signal, provided it had at least 10 counts and was more than double the signal of the second most abundant hashtag. Hashtag count data were also visualized using t-SNE to ensure clear separation of clusters corresponding to each hashtag. All single cells from the gene count matrix were uniquely matched to a single hashtag, thereby linking them unambiguously to their original sample.

#### Quality control (QC) and batch correction

Cells meeting any of the following criteria were excluded from the analysis: fewer than 500 RNA counts, dead cells with over 10% of counts mapping to mitochondrial genes, fewer than 500 TotalSeq-C counts, or positivity for two isotype controls (indicating non-specific TotalSeq-C antibody binding). Non-T cells were excluded based on the expression of the MNP gene signature, B cell signature, ILC gene signature, and the absence of T cell gene signature (score calculated using AddModuleScore_UCell)

#### Clustering and dimensionality reduction

Using Seurat v4.2.0 [https://pubmed.ncbi.nlm.nih.gov/31178118/], the variance-stabilizing transformation (VST method) was applied, and PCA was conducted on the top 2000 genes. Principal components explaining 80% of the total variance were selected for two-dimensional reduction using UMAP. Clustering was performed on these principal components using the FindClusters() function in Seurat.

#### Differential Expression

To determine differentially expressed genes, FindMarkers() within Seurat was used.

#### TCRαβ analyses

TCRαβ contigs for each single cell were obtained by aligning reads to reference genes (refdata-cellranger-vdj-GRCm38-alts-ensembl-7.0.0) using CellRanger software (v7.1.0) with default parameters. Immunarch (v1.0.0) was used to analyze TCRαβ repertoires. TCR gene usage and quasi-public clone neighborhood analyses were performed with TCRdist3(36) using Python (v3.10.10) and Jupyter notebook (v6.5.4).

### Spatial data processing

For the 10x Xenium spatial transcriptomics data, single cells from the day 3 and day 30 HDM samples were co-embedded with the Hurskainen et al. mouse lung single-cell RNA-seq reference (63) using scVI (66) and scANVI (62) after removing cells with < 10 transcript counts. The reference genes were subsetted to include only those in the Xenium panel. The scVI model was trained with n_layers=2, n_latent=30, and batch_key=’sample’ to distinguish the two Xenium samples and reference. The ‘CellType’ label in the Hurskainen et al. dataset was used for label prediction after consolidating labels with low cell counts in the reference or those not well separated by the genes in the Xenium panel. The reference was also filtered to remove cells with < 1000 counts to ensure only high quality cells were used for label prediction. The B cell, T cell, and DC labels were further refined by leiden subclustering following scANVI labeling. To label CD4 T cell populations consistently with the CITE-seq dataset, the Xenium and CITE-seq activated CD4 T cell populations were integrated using Seurat, subsetting each dataset to only the 185 genes robustly expressed in the CITE-seq CD4 T cells. Leiden clusters in the integrated embedding were labeled based on the CITE-seq cell identities.

CellCharter (64) was used to identify spatial neighborhoods, or zones, across multiple Xenium samples. Flu acute and memory timepoint samples were included in addition to the HDM samples, to facilitate identification of a robust TLS zone. To run CellCharter, an scVI model was first trained with n_layers=2, n_latent=30, and batch_key=’sample’. ScVI dimensions were then input to CellCharter (run with default parameters) for spatial neighborhood identification across a range of resolutions. The CellCharter neighborhoods were then consolidated based on the cell composition and spatial distribution. The adventitia and capsule zones were separated by assigning all cells within an average of 100 µm of the nearest 5 mesothelial cells to the capsule zone.

CD4 T cell colocalization analysis (Fig. 4F) on the Xenium dataset was performed using squidpy.gr.spatial_neighbors and squidpy.gr.interaction_matrix separately on activated CD4 T cells within each zone. Differential gene expression (DE) analysis on the Xenium dataset (Extended Data 7) was performed using scanpy.tl.rank_genes_groups with zones as groups. For the CD4 T cell DE test, genes were first filtered to include only those robustly expressed (expressed in at least 5% of cells on a per-subset basis) in both the CITE-seq and Xenium CD4 T cells. To identify gene expression gradients across spatial distances (Fig. 4G-H), the distance of all cells to TLS and bronchi zones were calculated. For the TLS, concave hulls were computed around the TLS zone and filtered by a minimum area threshold to distinguish mature TLS from more sparse groups of cells in the TLS zone. The distance to TLS for all cells was then computed as the distance to the closest TLS polygon, using the Python Shapely package. The distance to bronchi was computed as the average distance to the nearest 10 cells of the bronchi zone, Cells instide the TLS and cells in the bronchi zone were set to distance = 0 for the respective distance axis. For plotting along each distance axis, cells with distance > 450 µm were excluded. Gene expression gradients across each distance axis were visualized using scvelo.pl.heatmap. Radial distances outward from the TLS center and outward from the TLS edge in the surrounding 50 µm ring was calculated using the Shapely package (Figure 6), To quantify ligand-receptor interactions between CD4 T cells in the TLS core, TLS outer region, and TLS surroundings with all other cell types in these regions, CellChat was used in spatial mode (computeCommunProb with type = “truncatedMean”, trim = 0.1, interaction.range = 50, contact.range = 10).

### Xenium gene panel design

A custom 480 gene 10X Xenium panel was designed based on our previously published gene panel (10). Additionally, manually selected cytokine/cytokine receptor genes, immune related transcription factors, prostate-related genes, neuron-related genes, epithelium-related genes, stroma-related genes and type 2 immune cell-related genes were added to the existing list. Next, supervised and unsupervised PERSIST was run using as mouse intestine single-cell RNA reference dataset and a mouse lung scRNA dataset with the previously compiled set of genes as prior information, to yield a final list of 480 genes. The final gene list is included in Supplementary Table 1.

## Acknowledgments

This work was supported by NIH Award R01A1177874, P30DK0989507, CRI Lloyd J old STAR, Cancer Research Institute, NIH/NIAMS R01 AR080068

---

**Extended Data Figure 1:**
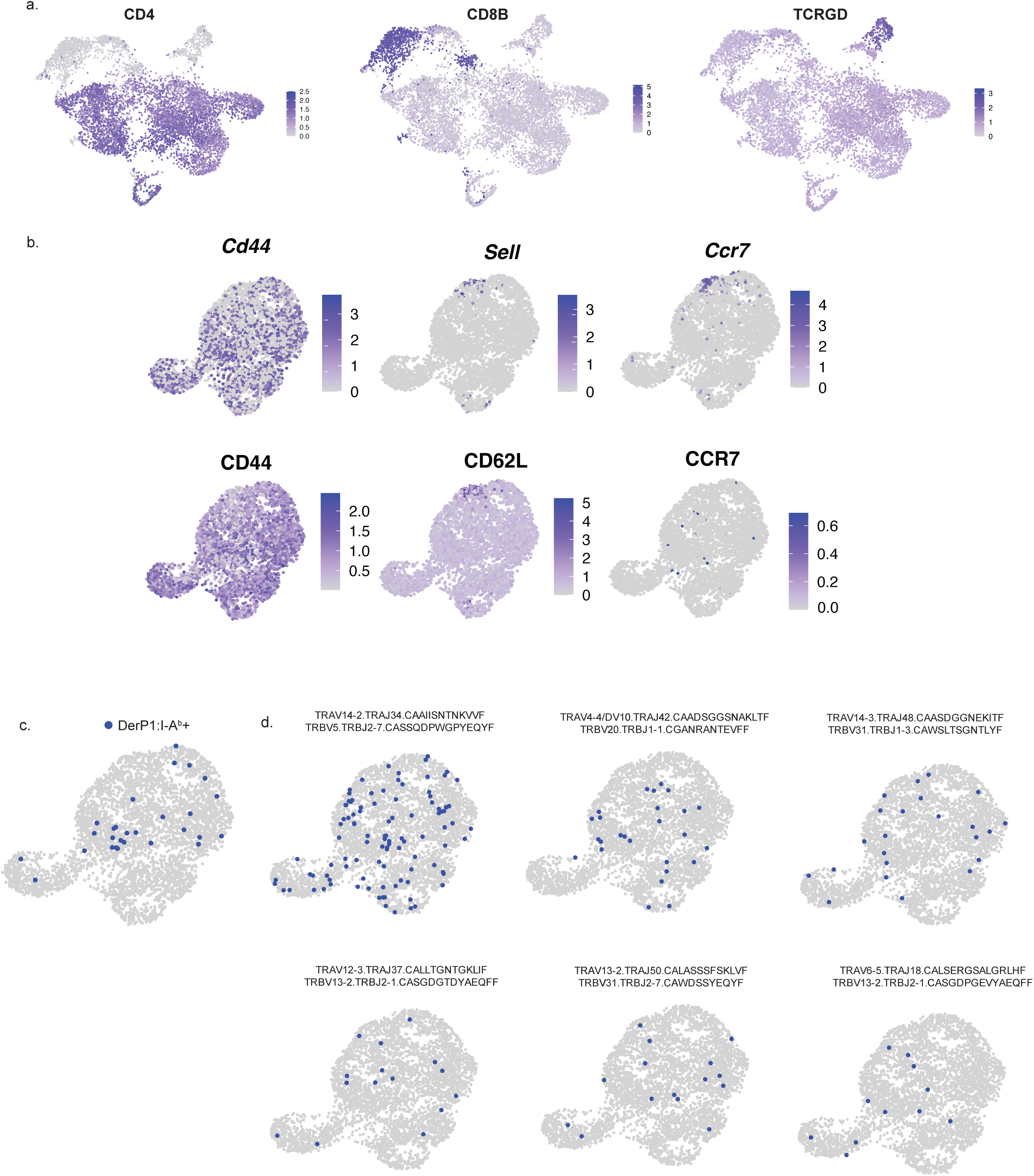
Additional profiling of T cells following HDM exposure. A) UMAPs showing the expression of CD4, CD8B, and TCRGD proteins among the combined single cell data set. B) UMAPs of lung CD4+ T cells at Secondary D3 and Secondary D30 showing the expression of the indicated gene or protein. C) UMAP highlighting Derp1:I-A^b^ tetramer cells D) UMAP highlighting expanded clonotypes. Each plot shows one clonotype expressing the indicated TCR alpha and TCR beta gene segments.

**Extended Data Figure 2:**
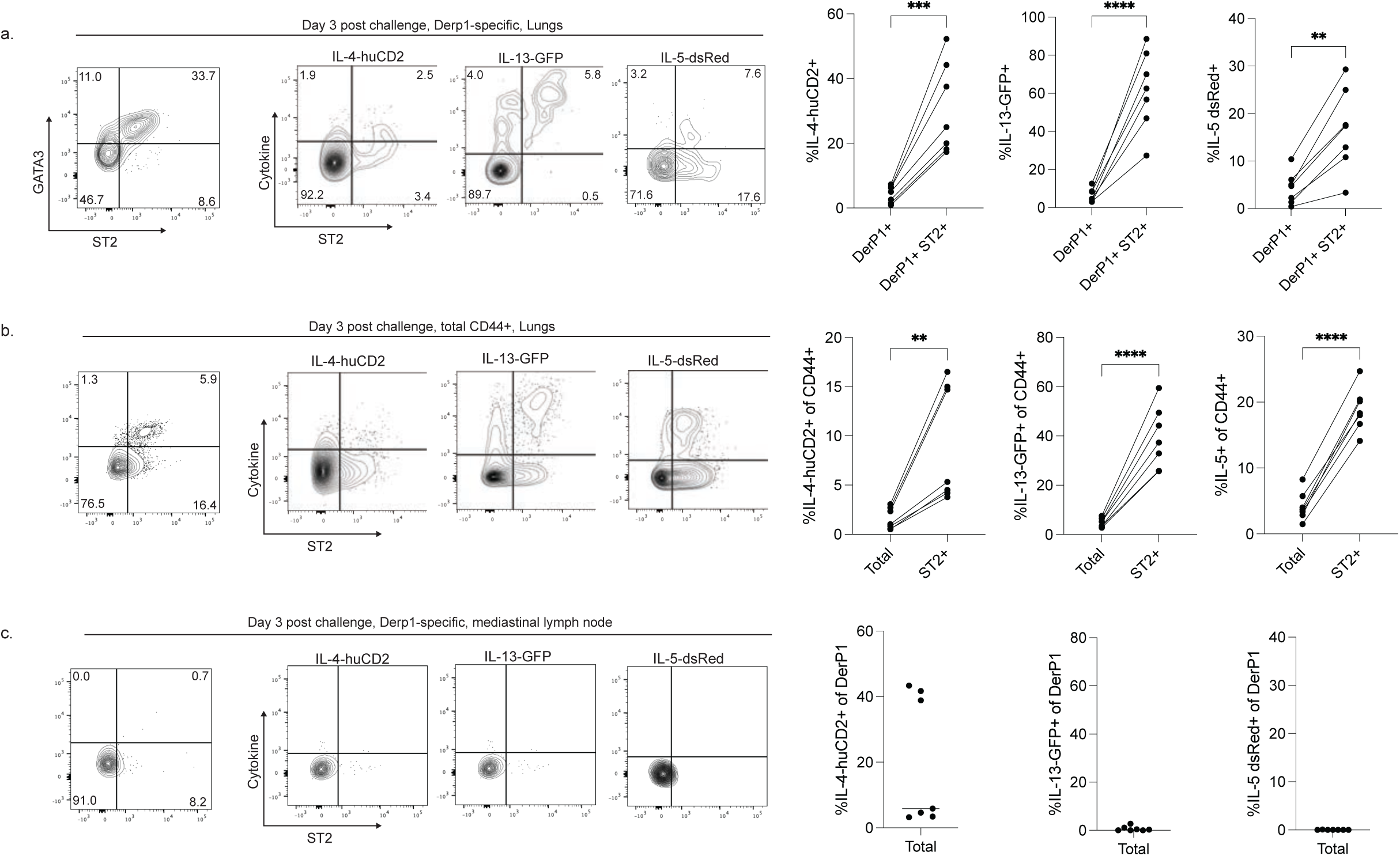
Additional data on cytokine production by CD4+ T cells. A-C) Flow cytometry plots and summary data showing expression of GATA3 as well as cytokine production by CD4+ T cells three days following HDM challenge. Flow plots are gated on Derp1-specific cells in the lungs (A), Derp1 tetramer-CD44+ cells in the lungs (B), and Derp1-specific cells in the mediastinal lymph node (C). Data are combined from two different experiments. Statistical significance determined using a paired t-test.

**Extended Data Figure 3:**
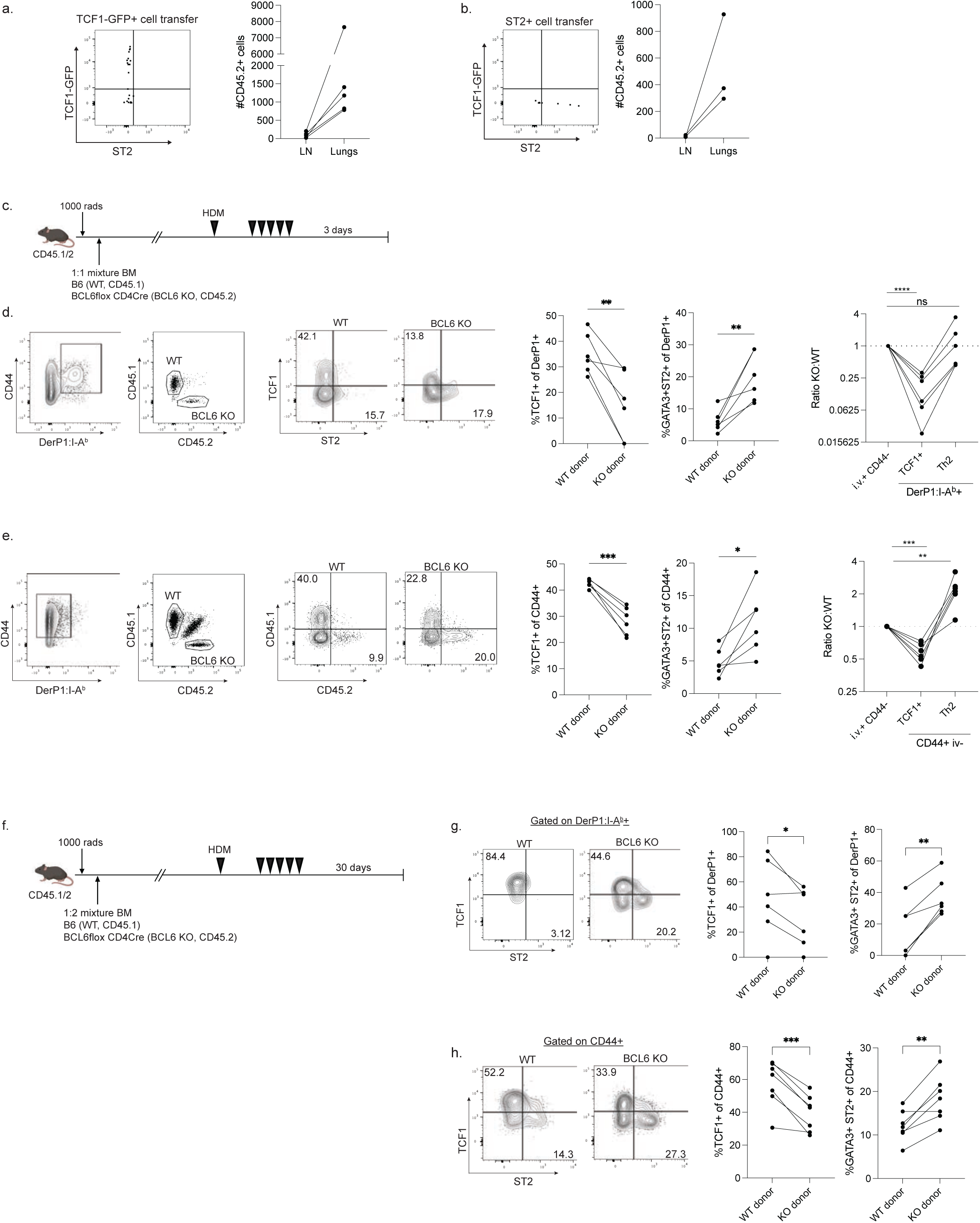
Extended data on adoptive transfer experiments and BCL6 KO mixed chimeras. A,B) Representative flow cytometry plots and summary data showing the phenotype and numbers of recovered TCF1-GFP+ cells (A) and ST2+ cells (B) from the mediastinal lymph node after adoptive transfer and restimulation, as outlined in Figure 2F. C) Experimental outline for BCL6-deficient mixed bone marrow chimeras. D,E) Representative flow plots and summary data showing the identification and phenotype of Derp1-specific lung CD4+ T cells (D) and total CD44+ (E) derived from the indicated bone marrow genotype. F) Experimental outline of memory BCL6-deficient mixed bone marrow chimeras. G,H) Flow cytometry plots and summary data of Derp1-specific (G) and total CD44+ CD4+ T cells (H) showing the phenotype of the cells generated from the indicated bone marrow donor. Data in D-H are combined from two independent experiments.

**Extended Data Figure 4:**
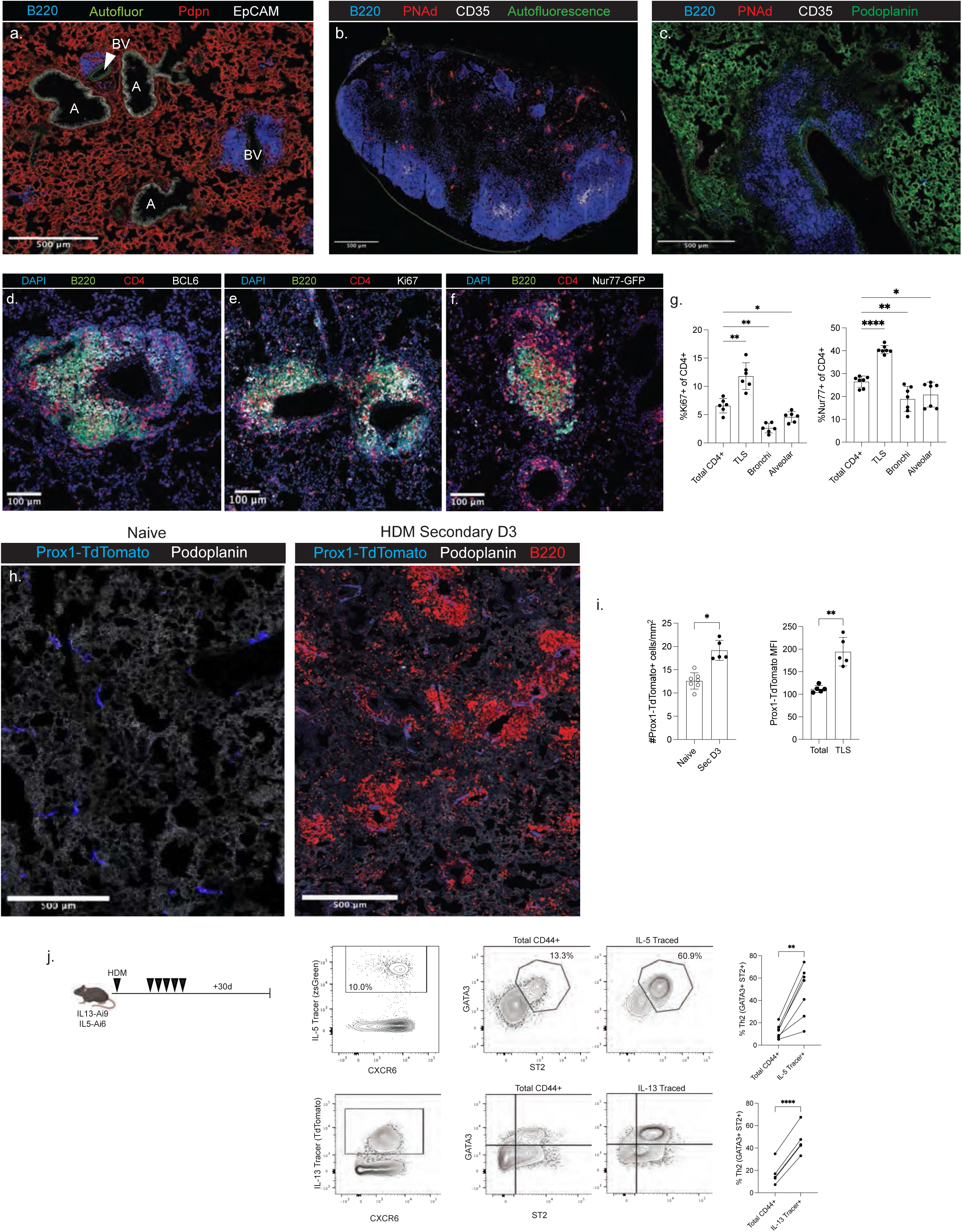
Characterization of lung TLS induced by HDM. A) Representative staining to identify anatomical sites in the lungs: bronchi (labeled as “A”, EpCAM+), alveoli (Podoplanin+), Blood Vessels (labeled as “BV”, Autofluorescence^hi^, EpCAM-), and tertiary lymphoid structures (TLS, dense B220+). B) Confocal microscopy of the mediastinal lymph node following HDM exposure showing B cell follicles (B220+), follicular dendritic cells (FDC, CD35+), and high endothelial venules (HEV, PNAd+). C) Representative staining of a lung TLS showing the lack of FDCs and HEVs. D) Representative staining of BCL6 in the lung TLS. E) Representative staining of Ki67. F) Representative staining of Nur77-GFP. G) Summary data showing the expression of Ki67 and Nur77-GFP by CD4+ T cells in the indicated region of the lung tissue three days post-HDM exposure. CD4+ cells were categorized as close to TLS (<50um), bronchi (<50um), or alveolar (>50um from TLS, >50um from bronchi, >50um from blood vessels). H) Representative confocal images and quantification of lymphatic vessels in naive (left) and HDM-exposed (right) lungs from Prox1-CreER^T2^-Ai9 mice. I) Summary data on the number of Prox1 vessels and mean fluorescence of Prox1 across the tissue. J) Flow data showing the detection and phenotype of IL-5- and IL-13-Traced cells in the lungs 31 days post-HDM exposure.

**Extended Data Figure 5:**
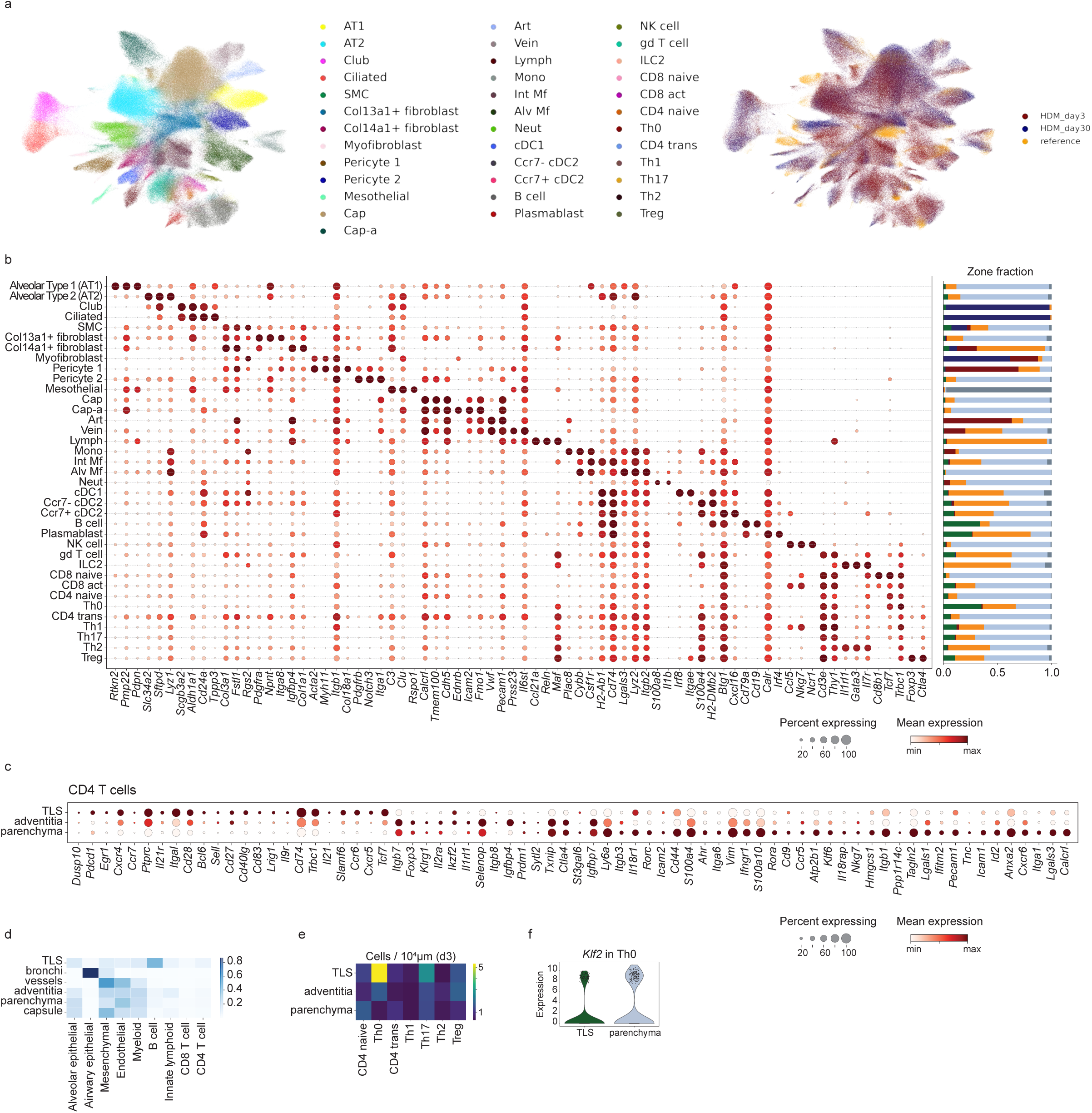
Additional characterization of lung niches with spatial transcriptomics. A) UMAP representation of Xenium lung data after scANVI integration with single-cell RNA-seq reference, colored by all labeled cell types (left, Xenium only) and Xenium sample or reference designation (right). B) Marker gene expression (left) and fraction in each zone (right) for labeled cell types. C) Differentially expressed genes (DEG) between major CD4-containing zones at day 3, with input genes filtered by expression in CITE-seq and Xenium data; genes included in heatmap have adjusted p value < 0.05 and abs(log_2_FC) > 1. D) Composition of each zone by major cell type, row scaled. E) Density of CD4 subsets in each major CD4-containing zone by zone tissue area, as in Figure 5E but additionally including naïve CD4 T cells. F) *Klf2* expression in Th0 labeled cells at day 3 in the TLS or parenchyma zones.

**Extended Data Figure 6:**
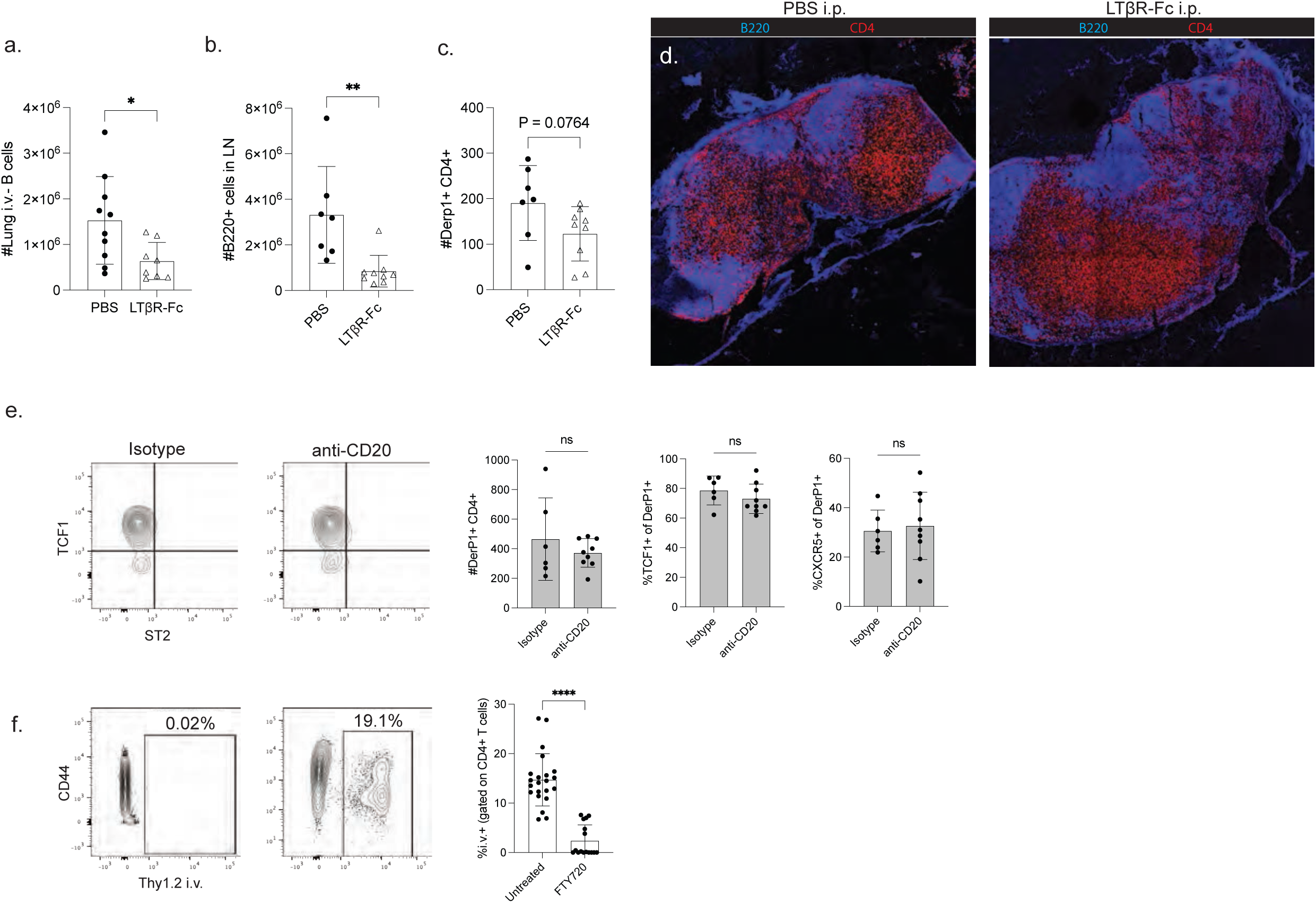
Additional data related to manipulations to TLS. A-C) Quantification of A) B cells in the lung tissue parenchyma, B) B cells in the draining lymph node, and C) Derp1-specific CD4 T cells in the draining lymph node of mice given the indicated treatments during HDM exposure as outlined in Figure 5A. D) Confocal microscopy of tissue sections from non-draining inguinal lymph nodes showing the bulk organization of the B cell follicle and T cell zone. Data in A-C are combined from two experiments, and data in D are representative of two experiments. Statistical significance determined with a t-test. E) Lymph node data from anti-CD20 treatment experiment (from Figure 5). Flow plots and summary data showing B cells and the phenotype of Derp1-specific CD4+ T cells in the mediastinal lymph node following lung B cell depletion. F) Representative flow plots and summary data of intravascular Thy1.2 antibody labeling of T cells in the lungs following FTY720 treatment. Data in F are pooled from 6 different experiments.

**Extended Data 7:**
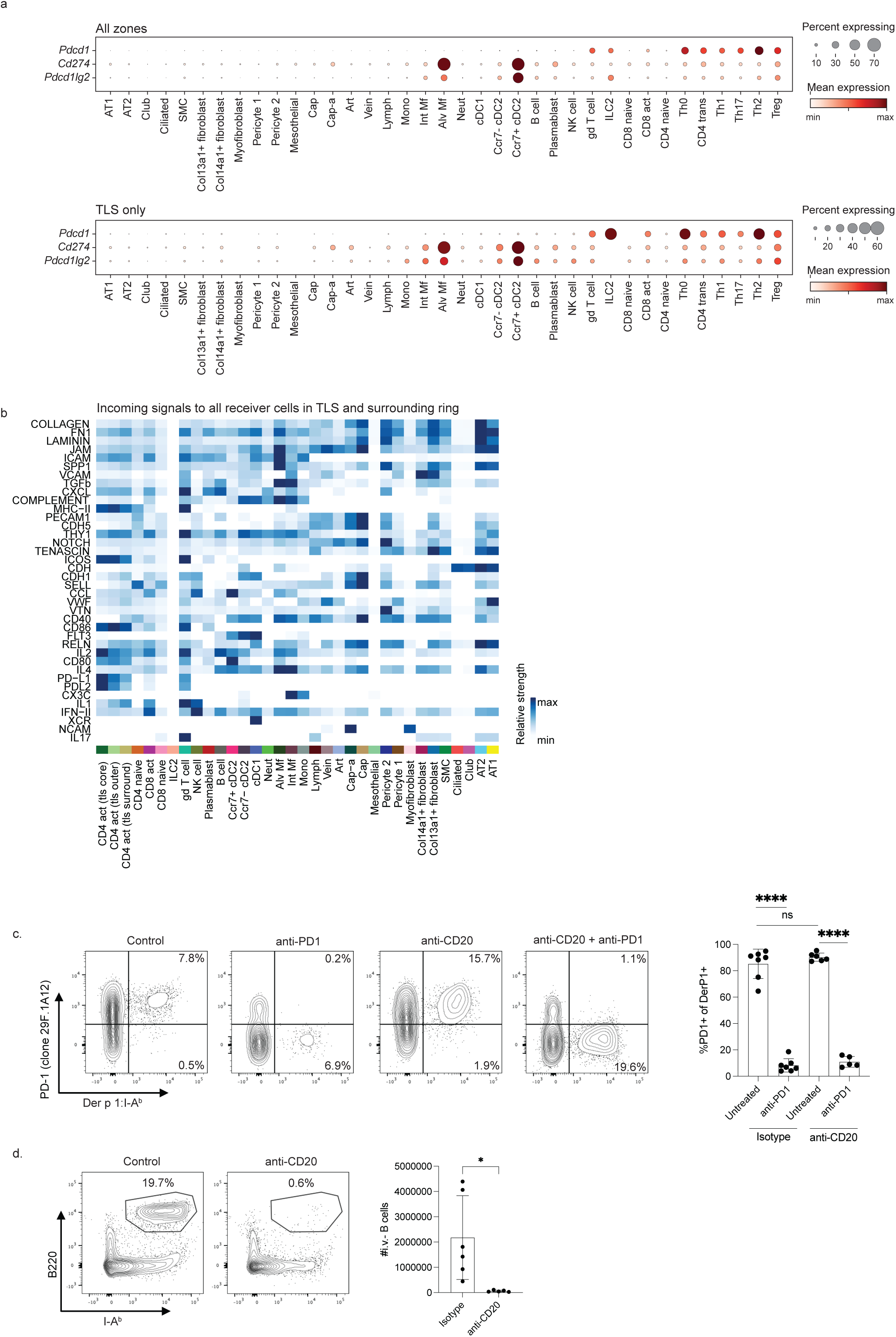
Additional data on PD1 ligand expression and PD1 blockade. A) *Pdcd1*, *Cd274*, and *Pdcd1lg2* expression by cell type in all zones (top) or specifically in the TLS zone (bottom). B) Incoming signaling pathways to all receiver cells in TLS and surrounding ring, scaled across signal strength was scaled across all receiver cells (rows). C) Representative flow cytometry plots and summary data showing PD1 expression by Derp1-specific T cells following the indicated treatment. D) Flow cytometry plots and summary data showing B cell quantification in mice that received anti-CD20 treatment or an isotype control. Data in C are combined from three experiments. Data in D are combined from two experiments. Statistical significance was determined using a t test.

## References

1. Gebhardt T, Wakim LM, Eidsmo L, Reading PC, Heath WR, Carbone FR. Memory T cells in nonlymphoid tissue that provide enhanced local immunity during infection with herpes simplex virus. Nat Immunol. 2009 May;10(5):524–30. doi: 10.1038/ni.1718. Epub 2009 Mar 22. PMID: 19305395.

2. Jiang X, Clark RA, Liu L, Wagers AJ, Fuhlbrigge RC, Kupper TS. Skin infection generates non-migratory memory CD8+ T(RM) cells providing global skin immunity. Nature. 2012 Feb 29;483(7388):227–31. doi: 10.1038/nature10851. PMID: 22388819; PMCID: PMC3437663.

3. Ariotti S, Hogenbirk MA, Dijkgraaf FE, Visser LL, Hoekstra ME, Song JY, Jacobs H, Haanen JB, Schumacher TN. T cell memory. Skin-resident memory CD8+ T cells trigger a state of tissue-wide pathogen alert. Science. 2014;346:101–105. doi: 10.1126/science.125480

4. Teijaro JR, Turner D, Pham Q, Wherry EJ, Lefrançois L, Farber DL. Cutting edge: Tissue-retentive lung memory CD4 T cells mediate optimal protection to respiratory virus infection. J Immunol. 2011;187:5510–5514. doi: 10.4049/jimmunol.1102243

5. Zundler, S. et al. Hobit- and Blimp-1-driven CD4+ tissue-resident memory T cells control chronic intestinal inflammation. Nat. Immunol. 20, 288–300 (2019).

6. Krebs, C. F. et al. Pathogen-induced tissue-resident memory TH17 (TRM17) cells amplify autoimmune kidney disease. Sci. Immunol. 5, eaba4163 (2020).

7. Hondowicz BD, An D, Schenkel JM, Kim KS, Steach HR, Krishnamurty AT, Keitany GJ, Garza EN, Fraser KA, Moon JJ, Altemeier WA, Masopust D, Pepper M. Interleukin-2-Dependent Allergen-Specific Tissue-Resident Memory Cells Drive Asthma. Immunity. 2016 Jan 19;44(1):155–166. doi: 10.1016/j.immuni.2015.11.004. Epub 2015 Dec 29. PMID: 26750312; PMCID: PMC4720536.

8. Rahimi RA, Nepal K, Cetinbas M, Sadreyev RI, Luster AD. Distinct functions of tissue-resident and circulating memory Th2 cells in allergic airway disease. J Exp Med. 2020 Sep 7;217(9):e20190865. doi: 10.1084/jem.20190865. PMID: 32579670; PMCID: PMC7478729.

9. Milner JJ, Toma C, He Z, Kurd NS, Nguyen QP, McDonald B, Quezada L, Widjaja CE, Witherden DA, Crowl JT, Shaw LA, Yeo GW, Chang JT, Omilusik KD, Goldrath AW. Heterogeneous Populations of Tissue-Resident CD8+ T Cells Are Generated in Response to Infection and Malignancy. Immunity. 2020 May 19;52(5):808–824.e7. doi: 10.1016/j.immuni.2020.04.007. PMID: 32433949; PMCID: PMC7784612.

10. Reina-Campos, M., Monell, A., Ferry, A. et al. Tissue-resident memory CD8 T cell diversity is spatiotemporally imprinted. Nature 639, 483–492 (2025). 10.1038/s41586-024-08466-x

11. Swarnalekha N, Schreiner D, Litzler LC, Iftikhar S, Kirchmeier D, Künzli M, Son YM, Sun J, Moreira EA, King CG. T resident helper cells promote humoral responses in the lung. Sci Immunol. 2021 Jan 8;6(55):eabb6808. doi: 10.1126/sciimmunol.abb6808. PMID: 33419790; PMCID: PMC8063390.

12. Son YM, Cheon IS, Wu Y, Li C, Wang Z, Gao X, Chen Y, Takahashi Y, Fu YX, Dent AL, Kaplan MH, Taylor JJ, Cui W, Sun J. Tissue-resident CD4+ T helper cells assist the development of protective respiratory B and CD8+ T cell memory responses. Sci Immunol. 2021 Jan 8;6(55):eabb6852. doi: 10.1126/sciimmunol.abb6852. PMID: 33419791; PMCID: PMC8056937.

13. Catarina Gago da Graça, Lisa G. M. van Baarsen, Reina E. Mebius; Tertiary Lymphoid Structures: Diversity in Their Development, Composition, and Role. J Immunol 15 January 2021; 206 (2): 273–281. 10.4049/jimmunol.2000873

14. Meylan M, Petitprez F, Becht E, Bougoüin A, Pupier G, Calvez A, Giglioli I, Verkarre V, Lacroix G, Verneau J, Sun CM, Laurent-Puig P, Vano YA, Elaïdi R, Méjean A, Sanchez-Salas R, Barret E, Cathelineau X, Oudard S, Reynaud CA, de Reyniès A, Sautès-Fridman C, Fridman WH. Tertiary lymphoid structures generate and propagate anti-tumor antibody-producing plasma cells in renal cell cancer. Immunity. 2022 Mar 8;55(3):527–541.e5. doi: 10.1016/j.immuni.2022.02.001. Epub 2022 Feb 28. PMID: 35231421.

15. Helmink BA, Reddy SM, Gao J, Zhang S, Basar R, Thakur R, Yizhak K, Sade-Feldman M, Blando J, Han G, Gopalakrishnan V, Xi Y, Zhao H, Amaria RN, Tawbi HA, Cogdill AP, Liu W, LeBleu VS, Kugeratski FG, Patel S, Davies MA, Hwu P, Lee JE, Gershenwald JE, Lucci A, Arora R, Woodman S, Keung EZ, Gaudreau PO, Reuben A, Spencer CN, Burton EM, Haydu LE, Lazar AJ, Zapassodi R, Hudgens CW, Ledesma DA, Ong S, Bailey M, Warren S, Rao D, Krijgsman O, Rozeman EA, Peeper D, Blank CU, Schumacher TN, Butterfield LH, Zelazowska MA, McBride KM, Kalluri R, Allison J, Petitprez F, Fridman WH, Sautès-Fridman C, Hacohen N, Rezvani K, Sharma P, Tetzlaff MT, Wang L, Wargo JA. B cells and tertiary lymphoid structures promote immunotherapy response. Nature. 2020 Jan;577(7791):549-555. doi: 10.1038/s41586-019-1922-8. Epub 2020 Jan 15. PMID: 31942075; PMCID: PMC8762581.

16. Iwata M, Sato A. Morphological and immunohistochemical studies of the lungs and bronchus-associated lymphoid tissue in a rat model of chronic pulmonary infection with Pseudomonas aeruginosa. Infect Immun. 1991 Apr;59(4):1514–20. doi: 10.1128/iai.59.4.1514-1520.1991. PMID: 2004830; PMCID: PMC257870.

17. GeurtsvanKessel CH, Willart MA, Bergen IM, van Rijt LS, Muskens F, Elewaut D, Osterhaus AD, Hendriks R, Rimmelzwaan GF, Lambrecht BN. Dendritic cells are crucial for maintenance of tertiary lymphoid structures in the lung of influenza virus-infected mice. J Exp Med. 2009 Oct 26;206(11):2339–49. doi: 10.1084/jem.20090410. Epub 2009 Sep 28. PMID: 19808255; PMCID: PMC2768850.

18. Richert LE, Harmsen AL, Rynda-Apple A, Wiley JA, Servid AE, Douglas T, Harmsen AG. Inducible bronchus-associated lymphoid tissue (iBALT) synergizes with local lymph nodes during antiviral CD4+ T cell responses. Lymphat Res Biol. 2013 Dec;11(4):196–202. doi: 10.1089/lrb.2013.0015. PMID: 24364842; PMCID: PMC3875184.

19. Sato Y, Jain A, Ohtsuki S, Okuyama H, Sturmlechner I, Takashima Y, Le KC, Bois MC, Berry GJ, Warrington KJ, Goronzy JJ, Weyand CM. Stem-like CD4+ T cells in perivascular tertiary lymphoid structures sustain autoimmune vasculitis. Sci Transl Med. 2023 Sep 6;15(712):eadh0380. doi: 10.1126/scitranslmed.adh0380. Epub 2023 Sep 6. PMID: 37672564; PMCID: PMC11131576.

20. Randolph, G.J. et al. Lymphoid Aggregates Remodel Lymphatic Collecting Vessels that Serve Mesenteric Lymph Nodes in Crohn Disease. Am J Pathol 186,3066–3073 (2016).

21. Rehal, S. & von der Weid, P.Y. TNFDeltaARE Mice Display Abnormal Lymphatics and Develop Tertiary Lymphoid Organs in the Mesentery. Am J Pathol 187,798–807 (2017).

22. Czepielewski, R.S. et al. Ileitis-associated tertiary lymphoid organs arise at lymphatic valves and impede mesenteric lymph flow in response to tumor necrosis factor. Immunity 54,2795–2811 e2799 (2021)

23. Elliot JG, Jensen CM, Mutavdzic S, Lamb JP, Carroll NG, James AL. Aggregations of lymphoid cells in the airways of nonsmokers, smokers, and subjects with asthma. Am J Respir Crit Care Med. 2004 Mar 15;169(6):712–8. doi: 10.1164/rccm.200308-1167OC. Epub 2004 Jan 7. PMID: 14711796.

24. Lee JJ, McGarry MP, Farmer SC, Denzler KL, Larson KA, Carrigan PE, et al. Interleukin-5 expression in the lung epithelium of transgenic mice leads to pulmonary changes pathognomonic of asthma. J Exp Med. (1997) 185:2143–56. doi: 10.1084/jem.185.12.2143

25. Hwang JY, Silva-Sanchez A, Carragher DM, Garcia-Hernandez ML, Rangel-Moreno J, Randall TD. Inducible Bronchus-Associated Lymphoid Tissue (iBALT) Attenuates Pulmonary Pathology in a Mouse Model of Allergic Airway Disease. Front Immunol. 2020 Sep 25;11:570661. doi: 10.3389/fimmu.2020.570661. PMID: 33101290; PMCID: PMC7545112.

26. Stern J, Pier J, Litonjua AA. Asthma epidemiology and risk factors. Semin Immunopathol. 2020 Feb;42(1):5–15. doi: 10.1007/s00281-020-00785-1. Epub 2020 Feb 4. PMID: 32020334.

27. Castillo JR, Peters SP, Busse WW. Asthma Exacerbations: Pathogenesis, Prevention, and Treatment. J Allergy Clin Immunol Pract. 2017 Jul-Aug;5(4):918–927. doi: 10.1016/j.jaip.2017.05.001. PMID: 28689842; PMCID: PMC5950727.

28. Van Dyken SJ, Nussbaum JC, Lee J, Molofsky AB, Liang HE, Pollack JL, Gate RE, Haliburton GE, Ye CJ, Marson A, Erle DJ, Locksley RM. A tissue checkpoint regulates type 2 immunity. Nat Immunol. 2016 Dec;17(12):1381–1387. doi: 10.1038/ni.3582. Epub 2016 Oct 17. PMID: 27749840; PMCID: PMC5275767.

29. Kratchmarov, R., Djeddi, S., Dunlap, G. et al. TCF1-LEF1 co-expression identifies a multipotent progenitor cell (TH2-MPP) across human allergic diseases. Nat Immunol 25, 902–915 (2024). 10.1038/s41590-024-01803-2

30. Moon JJ, Chu HH, Hataye J, Pagán AJ, Pepper M, McLachlan JB, Zell T, Jenkins MK. Tracking epitope-specific T cells. Nat Protoc. 2009;4(4):565–81. doi: 10.1038/nprot.2009.9. PMID: 19373228; PMCID: PMC3517879.

31. Endo Y, Hirahara K, Iinuma T, Shinoda K, Tumes DJ, Asou HK, Matsugae N, Obata-Ninomiya K, Yamamoto H, Motohashi S, Oboki K, Nakae S, Saito H, Okamoto Y, Nakayama T. The interleukin-33-p38 kinase axis confers memory T helper 2 cell pathogenicity in the airway. Immunity. 2015 Feb 17;42(2):294–308. doi: 10.1016/j.immuni.2015.01.016. PMID: 25692703.

32. Upadhyaya B, Yin Y, Hill BJ, Douek DC, Prussin C. Hierarchical IL-5 expression defines a subpopulation of highly differentiated human Th2 cells. J Immunol. 2011 Sep 15;187(6):3111–20. doi: 10.4049/jimmunol.1101283. Epub 2011 Aug 17. PMID: 21849680; PMCID: PMC3445433.

33. Wang JM, Rambaldi A, Biondi A, Chen ZG, Sanderson CJ, Mantovani A. Recombinant human interleukin 5 is a selective eosinophil chemoattractant. Eur J Immunol. 1989 Apr;19(4):701–5. doi: 10.1002/eji.1830190420. PMID: 2659368.

34. Scott G, Asrat S, Allinne J, Keat Lim W, Nagashima K, Birchard D, Srivatsan S, Ajithdoss DK, Oyejide A, Ben LH, Walls J, Le Floc’h A, Yancopoulos GD, Murphy AJ, Sleeman MA, Orengo JM. IL-4 and IL-13, not eosinophils, drive type 2 airway inflammation, remodeling and lung

35. Im SJ, Hashimoto M, Gerner MY, Lee J, Kissick HT, Burger MC, Shan Q, Hale JS, Lee J, Nasti TH, Sharpe AH, Freeman GJ, Germain RN, Nakaya HI, Xue HH, Ahmed R. Defining CD8+ T cells that provide the proliferative burst after PD-1 therapy. Nature. 2016 Sep 15;537(7620):417–421. doi: 10.1038/nature19330. Epub 2016 Aug 2. PMID: 27501248; PMCID: PMC5297183.

36. Bangs DJ, Tsitsiklis A, Steier Z, Chan SW, Kaminski J, Streets A, Yosef N, Robey EA. CXCR3 regulates stem and proliferative CD8+ T cells during chronic infection by promoting interactions with DCs in splenic bridging channels. Cell Rep. 2022 Jan 18;38(3):110266. doi: 10.1016/j.celrep.2021.110266. PMID: 35045305; PMCID: PMC8896093.

37. Xia Y, Sandor K, Pai JA, Daniel B, Raju S, Wu R, Hsiung S, Qi Y, Yangdon T, Okamoto M, Chou C, Hiam-Galvez KJ, Schreiber RD, Murphy KM, Satpathy AT, Egawa T. BCL6-dependent TCF-1+ progenitor cells maintain effector and helper CD4+ T cell responses to persistent antigen. Immunity. 2022 Jul 12;55(7):1200–1215.e6. doi: 10.1016/j.immuni.2022.05.003. Epub 2022 May 27. PMID: 35637103; PMCID: PMC10034764.

38. Wu T, Shin HM, Moseman EA, Ji Y, Huang B, Harly C, Sen JM, Berg LJ, Gattinoni L, McGavern DB, Schwartzberg PL. TCF1 Is Required for the T Follicular Helper Cell Response to Viral Infection. Cell Rep. 2015 Sep 29;12(12):2099–110. doi: 10.1016/j.celrep.2015.08.049. Epub 2015 Sep 10. PMID: 26365183; PMCID: PMC4591235.

39. Cardenas MA, Prokhnevska N, Sobierajska E, Gregorova P, Medina CB, Valanparambil RM, Greenwald R, DelBalzo L, Bilen MA, Joshi SS, Narayan VM, Master VA, Sanda MG, Kissick HT. Differentiation fate of a stem-like CD4 T cell controls immunity to cancer. Nature. 2024 Dec;636(8041):224–232. doi: 10.1038/s41586-024-08076-7. Epub 2024 Oct 23. Nature. 2024 Nov;635(8039):E9. doi: 10.1038/s41586-024-08303-1. PMID: 39443797.

40. Silva-Sanchez A, Randall TD. Role of iBALT in Respiratory Immunity. Curr Top Microbiol Immunol. 2020;426:21–43. doi: 10.1007/82_2019_191. PMID: 31974759; PMCID: PMC7875466.

41. Steele MM, Jaiswal A, Delclaux I, Dryg ID, Murugan D, Femel J, Son S, du Bois H, Hill C, Leachman SA, Chang YH, Coussens LM, Anandasabapathy N, Lund AW. T cell egress via lymphatic vessels is tuned by antigen encounter and limits tumor control. Nat Immunol. 2023 Apr;24(4):664–675. doi: 10.1038/s41590-023-01443-y. Epub 2023 Feb 27. Erratum in: Nat Immunol. 2023 Apr;24(4):729. doi: 10.1038/s41590-023-01491-4. PMID: 36849745; PMCID: PMC10998279

42. Dahlgren MW, Jones SW, Cautivo KM, Dubinin A, Ortiz-Carpena JF, Farhat S, Yu KS, Lee K, Wang C, Molofsky AV, Tward AD, Krummel MF, Peng T, Molofsky AB. Adventitial Stromal Cells Define Group 2 Innate Lymphoid Cell Tissue Niches. Immunity. 2019 Mar 19;50(3):707–722.e6. doi: 10.1016/j.immuni.2019.02.002. Epub 2019 Feb 26. PMID: 30824323; PMCID: PMC6553479.

43. Calvanese AL, Cecconi V, Stäheli S, Schnepf D, Nater M, Pereira P, Gschwend J, Heikenwälder M, Schneider C, Ludewig B, Silina K, van den Broek M. Sustained innate interferon is an essential inducer of tertiary lymphoid structures. Eur J Immunol. 2024 Oct;54(10):e2451207. doi: 10.1002/eji.202451207. Epub 2024 Jul 9. PMID: 38980268.

44. Harris P, Sridhar S, Peng R, Phillips JE, Cohn RG, Burns L, Woods J, Ramanujam M, Loubeau M, Tyagi G, Allard J, Burczynski M, Ravindran P, Cheng D, Bitter H, Fine JS, Bauer CM, Stevenson CS. Double-stranded RNA induces molecular and inflammatory signatures that are directly relevant to COPD. Mucosal Immunol. 2013 May;6(3):474–84. doi: 10.1038/mi.2012.86. Epub 2012 Sep 19. PMID: 22990623; PMCID: PMC3629368.

45. Hashimoto D, Chow A, Noizat C, Teo P, Beasley MB, Leboeuf M, Becker CD, See P, Price J, Lucas D, Greter M, Mortha A, Boyer SW, Forsberg EC, Tanaka M, van Rooijen N, García-Sastre A, Stanley ER, Ginhoux F, Frenette PS, Merad M. Tissue-resident macrophages self-maintain locally throughout adult life with minimal contribution from circulating monocytes. Immunity. 2013 Apr 18;38(4):792–804. doi: 10.1016/j.immuni.2013.04.004. PMID: 23601688; PMCID: PMC3853406.

46. Rodriguez AB, Peske JD, Woods AN, Leick KM, Mauldin IS, Meneveau MO, Young SJ, Lindsay RS, Melssen MM, Cyranowski S, Parriott G, Conaway MR, Fu YX, Slingluff CL Jr, Engelhard VH. Immune mechanisms orchestrate tertiary lymphoid structures in tumors via cancer-associated fibroblasts. Cell Rep. 2021 Jul 20;36(3):109422. doi: 10.1016/j.celrep.2021.109422. PMID: 34289373; PMCID: PMC8362934.

47. Son YM, Cheon IS, Li C, Sun J. Persistent B Cell-Derived MHC Class II Signaling Is Required for the Optimal Maintenance of Tissue-Resident Helper T Cells. Immunohorizons. 2024 Feb 1;8(2):163–171. doi: 10.4049/immunohorizons.2300093. PMID: 38345472; PMCID: PMC10916357.

48. Gräbner R, Lötzer K, Döpping S, Hildner M, Radke D, Beer M, Spanbroek R, Lippert B, Reardon CA, Getz GS, Fu YX, Hehlgans T, Mebius RE, van der Wall M, Kruspe D, Englert C, Lovas A, Hu D, Randolph GJ, Weih F, Habenicht AJ. Lymphotoxin beta receptor signaling promotes tertiary lymphoid organogenesis in the aorta adventitia of aged ApoE-/- mice. J Exp Med. 2009 Jan 16;206(1):233–48. doi: 10.1084/jem.20080752. Epub 2009 Jan 12. PMID: 19139167; PMCID: PMC2626665.

49. Conlon TM, John-Schuster G, Heide D, Pfister D, Lehmann M, Hu Y, Ertüz Z, Lopez MA, Ansari M, Strunz M, Mayr C, Angelidis I, Ciminieri C, Costa R, Kohlhepp MS, Guillot A, Günes G, Jeridi A, Funk MC, Beroshvili G, Prokosch S, Hetzer J, Verleden SE, Alsafadi H, Lindner M, Burgstaller G, Becker L, Irmler M, Dudek M, Janzen J, Goffin E, Gosens R, Knolle P, Pirotte B, Stoeger T, Beckers J, Wagner D, Singh I, Theis FJ, de Angelis MH, O’Connor T, Tacke F, Boutros M, Dejardin E, Eickelberg O, Schiller HB, Königshoff M, Heikenwalder M, Yildirim AO. Inhibition of LTβR signalling activates WNT-induced regeneration in lung. Nature. 2020 Dec;588(7836):151–156. doi: 10.1038/s41586-020-2882-8. Epub 2020 Nov 4. Erratum in: Nature. 2021 Jan;589(7842):E6. doi: 10.1038/s41586-020-03087-6. PMID: 33149305; PMCID: PMC7718297.

50. Jin S, Plikus MV, Nie Q. CellChat for systematic analysis of cell-cell communication from single-cell transcriptomics. Nat Protoc. 2025 Jan;20(1):180–219. doi: 10.1038/s41596-024-01045-4. Epub 2024 Sep 16. PMID: 39289562.Randolph G, Erlich E, Czepielewski R, Field R, Dunning T, Saleh L, Hoofnagle M, Tumanov A, Guilak F, Brestoff J. Distinct roles for LTalpha3 and LTalpha1beta2 produced by B cells contribute to their multi-faceted impact on ileitis. Res Sq [Preprint]. 2024 Feb 26:rs.3.rs-3962916. doi: 10.21203/rs.3.rs-3962916/v1. PMID: 38464070; PMCID: PMC10925464.

51. Randolph G, Erlich E, Czepielewski R, Field R, Dunning T, Saleh L, Hoofnagle M, Tumanov A, Guilak F, Brestoff J. Distinct roles for LTalpha3 and LTalpha1beta2 produced by B cells contribute to their multi-faceted impact on ileitis. Res Sq [Preprint]. 2024 Feb 26:rs.3.rs–3962916. doi: 10.21203/rs.3.rs-3962916/v1. PMID: 38464070; PMCID: PMC10925464.

52. Lai, Z., Kong, D., Li, Q. et al. Single-cell spatial transcriptomics of tertiary lymphoid organ-like structures in human atherosclerotic plaques. Nat Cardiovasc Res (2025). 10.1038/s44161-025-00639-9

53. Yu WW, Barrett JNP, Tong J, Lin MJ, Marohn M, Devlin JC, Herrera A, Remark J, Levine J, Liu PK, Fang V, Zellmer AM, Oldridge DA, Wherry EJ, Lin JR, Chen JY, Sorger P, Santagata S, Krueger JG, Ruggles KV, Wang F, Su C, Koralov SB, Wang J, Chiu ES, Lu CP. Skin immune-mesenchymal interplay within tertiary lymphoid structures promotes autoimmune pathogenesis in hidradenitis suppurativa. Immunity. 2024 Dec 10;57(12):2827–2842.e5. doi: 10.1016/j.immuni.2024.11.010. PMID: 39662091.

54. Singh M, Gautam N, Kaur M, Yadav D, Gupta P. Intravenous rituximab for the treatment of relapsing adult-onset asthma with periocular xanthogranuloma. Can J Ophthalmol. 2019 Jun;54(3):e115–e118. doi: 10.1016/j.jcjo.2018.08.013. Epub 2018 Nov 8. PMID: 31109495.

55. Casal Moura M, Berti A, Keogh KA, Volcheck GW, Specks U, Baqir M. Asthma control in eosinophilic granulomatosis with polyangiitis treated with rituximab. Clin Rheumatol. 2020 May;39(5):1581–1590. doi: 10.1007/s10067-019-04891-w. Epub 2020 Jan 2. PMID: 31897956.

56. Dasgupta, A., Radford, K., Arnold, D.M. et al. The effects of rituximab on serum IgE and BAFF. All Asth Clin Immun 9, 39 (2013). 10.1186/1710-1492-9-39

57. Siddiqui, Imran, et al. “Intratumoral Tcf1+ PD-1+ CD8+ T cells with stem-like properties promote tumor control in response to vaccination and checkpoint blockade immunotherapy.” Immunity 50.1 (2019): 195–211.

58. Kelli A. Connolly et al., A reservoir of stem-like CD8+ T cells in the tumor-draining lymph node preserves the ongoing antitumor immune response.Sci. Immunol.6,eabg7836(2021).DOI:10.1126/sciimmunol.abg7836

59. Sautès-Fridman, C., Petitprez, F., Calderaro, J. et al. Tertiary lymphoid structures in the era of cancer immunotherapy. Nat Rev Cancer 19, 307–325 (2019). 10.1038/s41568-019-0144-6

60. Anderson KG, Mayer-Barber K, Sung H, Beura L, James BR, Taylor JJ, Qunaj L, Griffith TS, Vezys V, Barber DL, Masopust D. Intravascular staining for discrimination of vascular and tissue leukocytes. Nat Protoc. 2014 Jan;9(1):209–22. doi: 10.1038/nprot.2014.005. Epub 2014 Jan 2. PMID: 24385150; PMCID: PMC4428344.

61. M. Stoeckius et al., Cell Hashing with barcoded antibodies enables multiplexing and doublet detection for single cell genomics. Genome Biol 19, 224 (2018).

62. Hurskainen M, Mizfkova I, Cook DP, Andersson N, Cyr-Depauw C, Lesage F, Helle E, Renesme L, Jankov RP, Heikinheimo M, Vanderhyden BC, Thébaud B. Single cell transcriptomic analysis of murine lung development on hyperoxia-induced damage. Nat Commun. 2021 Mar 10;12(1):1565. doi: 10.1038/s41467-021-21865-2. PMID: 33692365; PMCID: PMC7946947.

63. Lopez R, Regier J, Cole MB, Jordan MI, Yosef N. Deep generative modeling for single-cell transcriptomics. Nat Methods. 2018 Dec;15(12):1053–1058. doi: 10.1038/s41592-018-0229-2. Epub 2018 Nov 30. PMID: 30504886; PMCID: PMC6289068.

64. Xu C, Lopez R, Mehlman E, Regier J, Jordan MI, Yosef N. Probabilistic harmonization and annotation of single-cell transcriptomics data with deep generative models. Mol Syst Biol. 2021 Jan;17(1):e9620. doi: 10.15252/msb.20209620. PMID: 33491336; PMCID: PMC7829634.

65. Varrone M, Tavernari D, Santamaria-Martínez A, Walsh LA, Ciriello G. CellCharter reveals spatial cell niches associated with tissue remodeling and cell plasticity. Nat Genet. 2024 Jan;56(1):74–84. doi: 10.1038/s41588-023-01588-4. Epub 2023 Dec 8. PMID: 38066188.

